# Exploring the genome-wide expression level of the bacterial strain belonging to *Bacillus safensis* (MM19) against *Phomopsis viticola*

**DOI:** 10.1101/2024.05.03.592302

**Authors:** Ragıp Soner Silme, Ömür Baysal, Ahmet Can, Yiğit Kürüm, Ahmet Korkut, Kevser Kübra Kırboga, Agit Çetinkaya

## Abstract

**Introduction:** Rhizobacteria has the suppression ability to compete with pathogenic microorganisms and help for plant immunity and defense mechanism. Their growth and survival in rhizosphere ensure biological balance in favor of the host plant.

**Methods:** In our study, with agar diffusion assays, we found a rhizobacterium species belonging to *Bacillus safensis*, which can significantly suppress *Phomopsis viticola*. To elucidate the antagonistic mechanism, genome-wide gene expression profiling of *B. safensis* (strain MM19) was performed in the presence and absence of *P. viticola*. We used RNA-seq analysis to obtain a comprehensive overview of the responsive *B. safensis* whole gene expression classified according to biological and metabolic process to *P. viticola* concomitant growth in liquid culture.

**Results:** The differential gene expression profiles of *B. safensis* (MM19) revealed significantly increased expression of prominent genes related to thiamine biosynthesis involving various metabolites and enzymes that play role in the suppression of mycelium growth and pathogen inhibition. Correspondingly, the expression of three major genes (HOG1, FUS3, SGI) involving in the virulence of *P. viticola* was followed using qPCR analysis. HOG1 was the highest expressed gene in the pathogen when it co-cultivated with MM19. Based on these findings, we performed molecular docking and dynamics analysis to explore the interaction between HOG1 and thiamine, besides expression of network analysis constructed using Cytoscape.

**Discussion:** The results proved that the functional genomic data related to thiamine biosynthesis and corresponding pathways ensure a priming role in the antagonistic behavior of *B. safensis* (MM19) against *P. viticola* as a supporter for plant immunity.

## 1 Introduction

Biocontrol is an eco-friendly approach involving the promotion of plant growth, production of hormones or indirect suppression of plant pathogens, and induction of plant resistance. Rhizobacteria interact with plants through a range of mechanisms that influence their growth and health. The direct interactions between rhizobacteria and plant pathogens encompass a multifaceted relationship involving elements of competition and antibiosis. These interactions involve the production of various compounds, including antibiotics, cyclic lipoproteins, antimicrobial volatiles, siderophores, and exoenzymes, which collectively contribute to the dynamic interplay between these microorganisms (Raaijmakers et al., 2002; McSpadden Gardener, 2004).

*Bacillus* species have demonstrated the ability to mitigate disease severity in various economically significant crop species. *Bacillus* spp. are ubiquitous bacteria found in soils that show antagonistic properties against crop pathogens (Choudhary and Johri, 2009). Some strains secrete antibiotics that can directly inhibit the growth of fungal, oomycete, and bacterial plant pathogens (Toure et al., 2004). The genetic traits underlying biocontrol efficiency and regulation that play a role in the suppression of plant pathogen growth have been studied on these bacterial strains (McSpadden Gardener, 2004) on account of their ability to produce cell wall-degrading enzymes and antibiotic compounds (Baysal et al., 2008; 2013; Pandin et al., 2017). Some other strains of *Bacillus subtilis* also suppress diseases by inducing host defense mechanisms (Miljaković et al., 2020). However, additional functional studies are required to understand the ongoing mode of action related to biocontrol. Whereas genome sequences provide an insight to understand the role of traits and their importance with the studies carried out on mutant individuals. In view of gene regulation and subsequent activity, transcriptome analysis is an important tool for combining the effects of different traits. To the best of our knowledge, these kinds of studies are few and many available biocontrol agent and pathogen interactions remain unclear.

*Bacillus safensis* is a Gram-positive and aerobic chemoheterotroph with spore-forming rod bacterium showing resistant ranges in temperature between 30 and 37°C (Satomi et al., 2006). It is also resistant to 10% salinity and pH level of 5-6. In a previous study, a strain of *B. safensis* has shown resistance to UV radiation and hydrogen peroxide (Tirumalai et al., 2013). *B. safensis* has the potential to enhance plant growth, both through direct and indirect mechanisms. This includes the inhibition of fungal pathogens, and the reduction of salt-induced stress in plants (Chakraborty et al., 2018). Wu et al. (2019) found that *B. safensis* strain ZY16 is an endophytic bacterium that can degrade hydrocarbons, produce biosurfactants, tolerate salt, and promote plant growth through IAA production, siderophore synthesis, and phosphate solubilizing activity.

*Phomopsis viticola* belongs to the serious pathogens of grapevine cultivated in many regions of the world and is known as Phomopsis dead arm disease (Scheper et al., 1997). Recent research findings have indicated that pruning wounds remain vulnerable to infection for an extended period, with susceptibility persisting for a minimum of three weeks following the pruning process (van Niekerk et al., 2011). One of the strategies for grapevine protection against this pathogen is to reduce *P. viticola* inoculum on canes, both during the vegetative and dormant periods, which also results in protection against fungus distribution (Castillo-Pando et al., 1997). Therefore, chemical applications are traditionally recommended (Ramsdell, 1995).

In the management of Phomopsis dead arm disease, a range of early-season spray treatments, including copper oxychloride, copper oxychloride/sulphur, copper sulphate/lime, folpet, fosetyl-Al + mancozeb, probineb, sulphur and strobilurin-based fungicides, have been sprayed (Mostert and Crous, 2000). There are no fungicides available that are both effective and environment-friendly in suppressing the fungus. Furthermore, the research conducted by Munkvold and Marois (1993) revealed that the efficacy of chemicals such as benomyl exhibited a decline in effectiveness two weeks after application. These findings have shown the importance of finding alternative methods that could replace protective pruning and chemical application. There are no data on the biocontrol of *P. viticola* by *B. safensis* treatment. Depending on the above-mentioned situation, the importance of a rhizobacterium that can control Phomopsis in terms of biological control becomes evident.

Whole transcriptome analysis with RNA-seq has opened a new dimension to understand the gene regulation and how it changes on account of the interactions between the host and the pathogen, which can shed light on the mechanisms of pathogenicity, host defense, and their interplay in various conditions (Wolf et al., 2018). Currently, RNA-seq is the major method used in transcriptomics studies and RNA plays a key role in various cellular processes. Therefore, investigation of the identity, function, and abundance of transcribed RNA molecules (i.e., transcripts) are crucial for understanding cellular behavior. Advances in the field of RNA biology were mainly driven by the development of novel technologies and methods that allow researchers to study different aspects of transcripts in an increasingly efficient manner. Therefore, we investigated the biocontrol agent and pathogen interaction using the RNA-seq technique (Figure 1 & 2).

**FIGURE 1.**
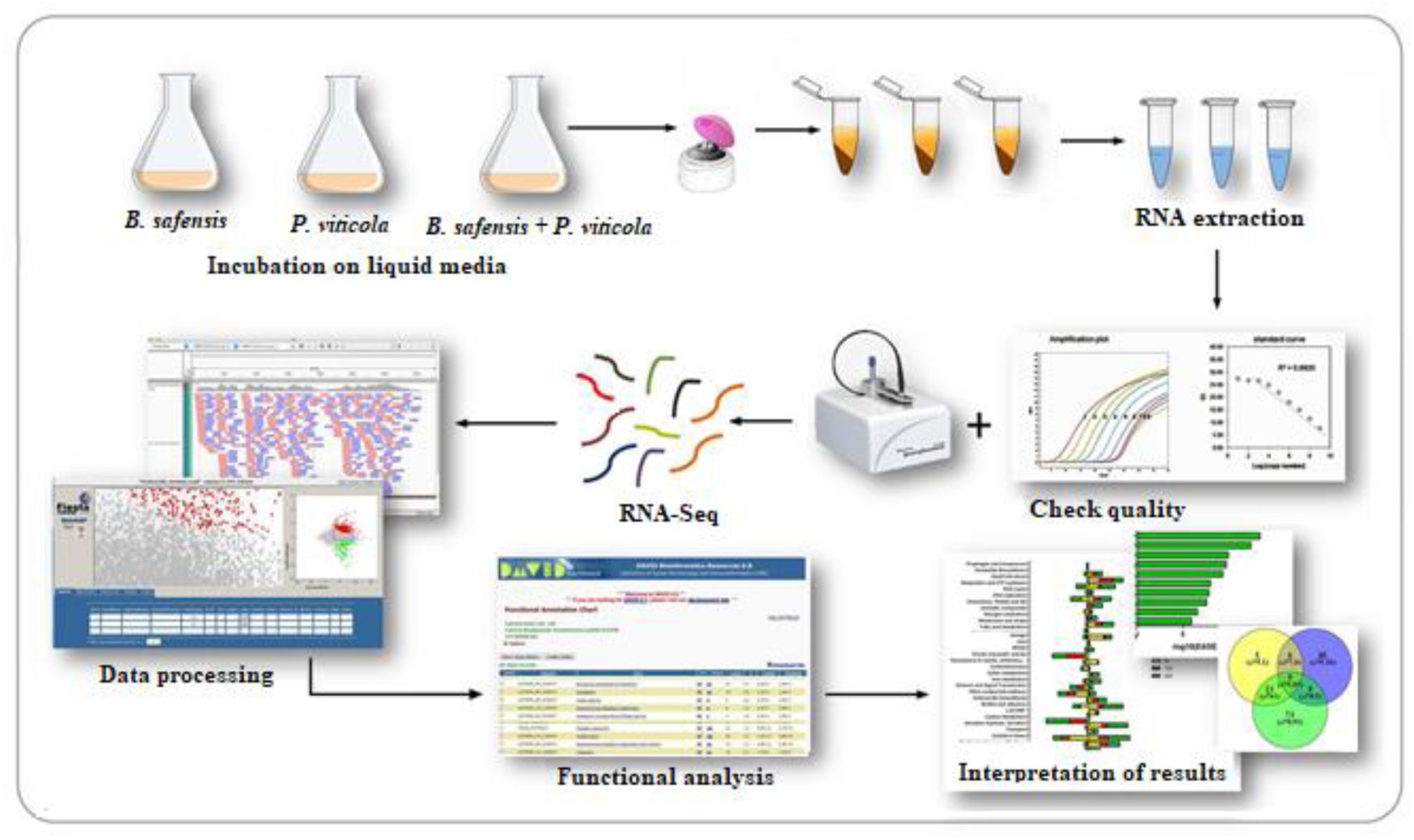
Workflow of RNA-seq studies. Workflow of the general procedure explained in these protocols. The main wet and *in silico* steps are summarized in the figure.

**FIGURE 2.**
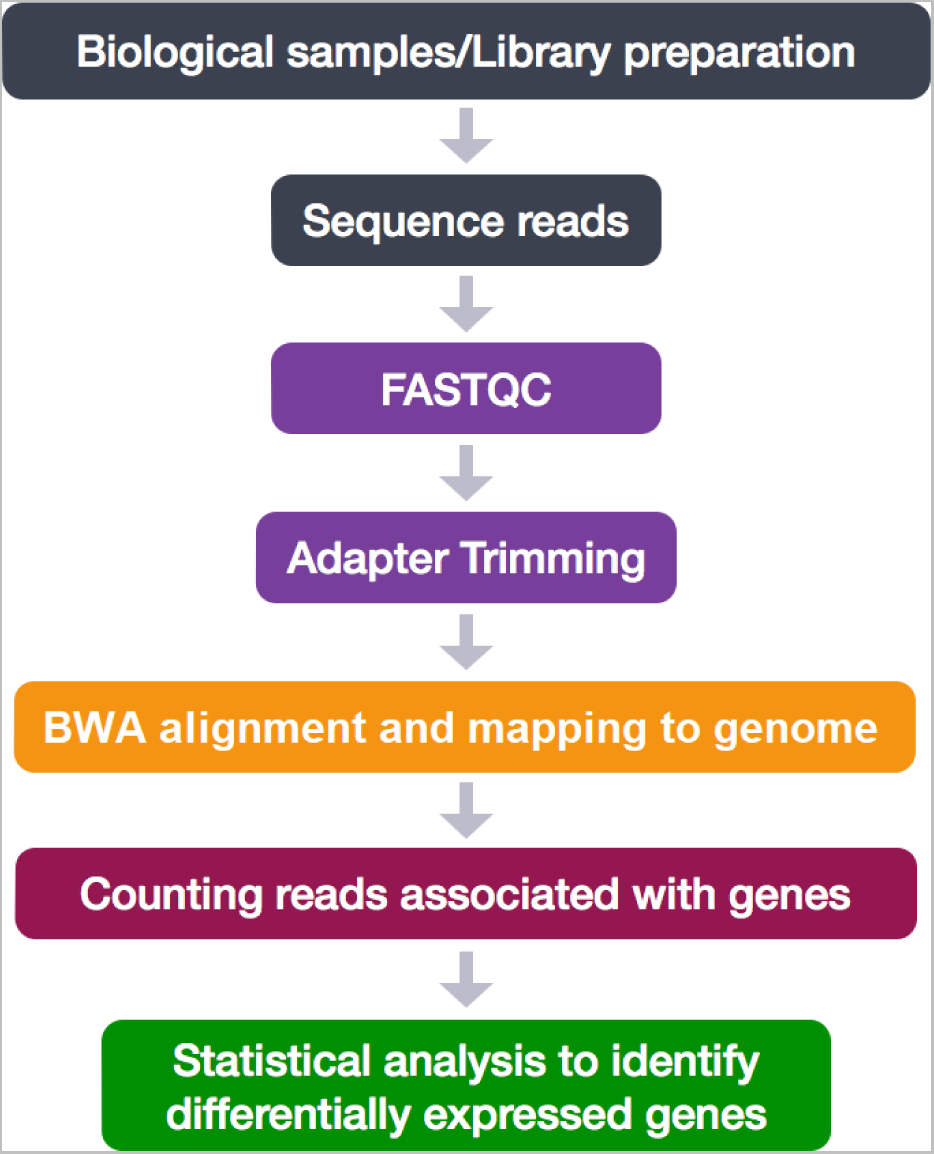
Overview of the work-chart for assembly of RNA-seq reads for B. *safensis* MM19.

This study aimed to investigate the alterations in the transcriptional responses exhibited by the bacterial strain MM19 against *P. viticola*. We conducted a comprehensive analysis of the genome-wide gene expression patterns in *B. safensis* in the presence and absence of *P. viticola*. This investigation was performed to elucidate the microbial responses of *B. safensis* when coexisting with *P. viticola* within *in vitro* conditions.

## 2 Materials and Methods

### 2.1 Microorganisms and culture conditions

The bacterial strain *Bacillus safensis* MM19 (Molecular Microbiology lab, Muğla, 2019) was isolated from soil samples of crude petroleum pollutants contaminated seashore near petroleum refinements (Aliaga, İzmir, Turkey). The bacterium was cultured on nutrient broth agar (NBA, Merck) and preserved at -20°C. The morphological characteristics, Gram’s reaction, spore formation and motility of the bacterial strain MM19 were examined by microscopic observations. The carbon source preferences of the strain were determined with Biolog GENIII MicroPlate assay according to the manufacturers’ directions by Dr. Küsek (Sütçü İmam University/Kahramanmaraş).

*P. viticola* samples with diseased grape seedling origin were obtained from the Bornova Plant Protection Research Institute/Izmir. The pathogen was previously identified at the species and genus level through ITS gene sequence analyses. Subsequently, the pathogen was cultured on potato dextrose agar (PDA, Merck) medium at 20°C.

### 2.2 Antagonistic effect of MM19 using single layer assay

*In vitro* experiment, to determine the antibiosis effect, the assay was conducted using single-layer methods. Fungal sections excised from 5-day-old *P. viticola* cultures were placed on a fresh PDA medium. When the pathogen mycelium reached at the edge of the Petri plates, a sample of the fungi near the center was excised using a cork borer. *P. viticola* culture was then centrally transferred to a culture medium consisting of PDA (pH 6.2), then the bacterium was inoculated as positioned two streaks with 2 cm distance reciprocally at either side of the mycelial plug. Later, the Petri dishes were incubated at 23°C for one month, the growth of mycelium was compared to be six replicates involving control groups in which only *P. viticola* was present.

The growth of mycelium was measured at 7, 10 and 12 days post-inoculation (dpi) compared to the control groups. Radius measurements were obtained from Petri dishes representing fungal growth in control samples and bacteria inoculated (treated groups). The data obtained were subjected to Student’s t-test analysis considering Bootstrap confidence level using SPSS (IBM SPSS Statistics for Windows, Version 22.0. Armonk, NY: IBM Corp.).

### 2.3 Reciprocally growth of the microorganisms in liquid culture to follow interaction and RNA isolation

For RNA isolation, the optical density (OD) of *B. safensis* depending on colony growth was measured *in vitro* conditions as single (control) or dual liquid culture systems at which *B. safensis* replaced alone and interacting with *P. viticola*. To determine whole expressed genes playing role in antibiosis, the liquid culture was consisted of equal amounts (50/50%) of potato dextrose (PD) and nutrient broth (NB) medium. The fungi obtained from 5-day-old *P. viticola* cultures were excised from PDA medium using a cork borer and released into 15-ml glass bottles containing PD liquid culture. Concomitantly, bacteria were inoculated into liquid culture and growth until the OD level reached 1.0. Then, two different inoculated liquid cultures containing bacteria and fungi were mixed into a 30 ml Erlenmayer flask and performed until the common OD level reached up to 1.0, of which shaking culture needs 3.5 h. Then, the RNA extraction procedure given below was followed.

During RNA extraction, the bacterial solution was filtered using sterile Whatman paper to separate fungal mycelium. Then, the bacterial solution was gently rinsed with the fungicide penconazole (10 µl / 15 ml) (100 g/l CAS No: 66246-88-6) for maximum 2 min, to minimize contamination with fungi in bacterial RNA.

### 2.4 Equations Bacterial RNA extraction and sample preparation for RNA-seq analysis

Harvested bacterial cells treated with fungicide were subjected to RNA extraction using a GeneJET RNA Purification Kit (K0372, Thermofischer) according to the manufacturer’s instructions for extraction of total RNA from bacterial cells. Total RNA was quantified using an ND-1000 spectrophotometer (NanoDrop Technologies, Delaware, USA).

Total RNA served as the input material for the preparation of RNA samples. Sequencing libraries were constructed using the NEBNext Ultra RNA Library Preparation Kit for Illumina (NEB, USA, Catalog #: E7530L) in accordance with the manufacturer’s guidelines. Index codes were introduced to associate individual sequences with their respective samples. mRNA was isolated from total RNA using poly-T oligo-affixed magnetic beads. The fragmentation process was conducted by subjecting the RNA to elevated temperatures in the presence of divalent cations and performed within the NEB Next First Strand Synthesis Reaction Buffer (5X). The first strand of complementary DNA (cDNA) was synthesized using a random hexamer primer and M-MuLV reverse transcriptase, which possesses ribonuclease H (RNase H) activity. The synthesis of the second strand of complementary DNA (cDNA) was performed using DNA polymerase I in conjunction with RNase H. The remaining overhang regions were converted into blunt ends by the enzymatic activities of exonuclease and polymerase. After the adenylation of the 3’ ends of DNA fragments, NEB Next adaptors featuring a hairpin loop structure were ligated to facilitate the preparation for hybridization. To selectively isolate cDNA fragments ranging from 370 to 420 base pairs in length, the library fragments were purified using the AMPure XP system (Beverly, USA). Subsequently, 3 µL volume of USER Enzyme (NEB, USA) was applied to the size-selected, adaptor-ligated cDNA, incubated at 37°C for 15 min, and then subjected to a 5-min incubation at 95°C before PCR amplification. Following these steps, polymerase chain reaction (PCR) was performed using Phusion High-Fidelity DNA polymerase, along with Universal PCR primers and Index (X) Primer. Then, the PCR products were prepared using the AMPure XP system, and the quality of the library was evaluated using the Agilent 5400 system (Agilent, USA). Quantification of the library was achieved through quantitative polymerase chain reaction (QPCR), resulting in a concentration of 1.5 nM. The whole validated libraries were combined and subjected to sequencing on Illumina platforms belonging to Ficus Bio Co. (Ankara/ Turkey), employing paired-end 150 base pair (PE150) sequencing, specifically on the Illumina HiSeq 2500 system. Sequencing depth was determined based on effective library concentration and requisite data quantity.

### 2.5 Raw data and bioinformatics analysis

The original fluorescence image files obtained from the Illumina platform are transformed to short reads (raw data) by base calling, and these short reads are recorded in FASTQ format (Chen et al., 2018), which contains sequence information and corresponding sequencing quality information. Fastp (version 0.23.1) was used to perform basic statistics on the quality of the raw reads (Chen et al., 2018). The steps of data processing involve a) discarding paired reads if either one read contains adapter contamination, b) discarding paired reads if more than 10% of bases are uncertain in either one read and c) discarding paired reads if the proportion of low quality (Phred quality <5) bases is over 50% in either one read.

Sequence artifacts, including reads containing adapter contamination, low-quality nucleotides, and unrecognizable nucleotide (N), undoubtedly set the barrier for subsequent reliable bioinformatics analysis. The paired-end reads for each sample were imported into the Galaxy software platform (The Galaxy Community, 2022) and mapped to the reference genome *Bacillus safensis* (https://www.ncbi.nlm.nih.gov/datasets/genome/GCF_003254445.1/) using default settings, except for the maximum insert size of the paired-end library. FastQ data were subjected to optimized trimmomatic tools (adjusted parameters for MINLEN, LEADING, CROP and HEADCROP) to remove adaptor sequences, for fastQC analysis. Then, RNA STAR was used to obtain BAM files from single- and pair-end sequences after MultiQC analysis and then visualized using Tablet (Hutton Institute ver. 1.21.02.08) (Milne et al., 2013). The multiple count data sets for multiple samples were retrieved and converted to a single file using feature counts. Then, the data counts were subjected to DeSeq2 analysis using Phyton (ver. 2023.2 Pycharm) through Sambomics tools to determine upregulated and downregulated genes and logFc data. The data were submitted to the IDEP platform to visualize the heatmap and other options related to gene counts (Ge et al., 2018). To determine WP values corresponding to gene ID, BLAST analysis was also performed. To annotate the expressed genes, BLASTx searches (E value < 10-3) were performed between nr and unigenes. BLAST results were imported to the Blast2GO program for further annotation of the unigenes (Conesa et al., 2005). Genes were attributed as differentially expressed if the fold change was greater than ±2 and with an FDR corrected p-value of ≤ 0.05.

Gene enrichment analysis was carried out with ShinyGO v0.741 (Ge et al., 2020) based on *Bacillus subtilis* STRINGdb and then visualized using SRTplot (Simple Real-Time Plotter, 2013). Because there was no available specific genomic data available for *B. safensis* in these analyses tools, we also compared the whole protein expression mapping obtained by the UNIPROT database considering the available data for *B. subtilis*.

### 2.6 Enrichment gene interaction map and network analysis

An enrichment map was generated through the utilization of files derived from RNA-seq output using Cytoscape, an open-source and freely accessible plugin designed for network analysis, specifically utilized for the visualization of enriched pathways within a network context. This process involves incorporating elements such as expression profiles, hallmark genesets, enrichment sets, and the specification of parameters such as p-value and overlap coefficient (Reimand et al., 2019). The enrichment map visually represents pathways as a network, wherein nodes correspond to distinct pathways and are interconnected by edges indicating shared common genes within the respective pathways. Moreover, the examination of individual gene interactions pertaining to diverse pathways involves the selection of relevant genes, and subsequently, a network is constructed by annotating these genes through the String Database, utilizing Cytoscape software (Merico et al., 2011).

### 2.7 qRT-PCR analysis of three selected major genes responsible for virulence

Fungal solution was treated with 10 μl Ampicillin (1000 mg/ml) for 15 min to remove bacterial cells. Harvested fungal mycelium from the liquid culture was used for total RNA extraction by GeneJET RNA Purification Kit (K0372, Thermofischer) according to the manufacturer’s instructions for yeast protocol. Total RNA was measured using spectrophotometry and used for qRT-PCR analysis.

Due to limited studies on specific virulence genes of *P. viticola*, we carried out qRT-PCR analysis from extracted RNA of fungi, which was concomitantly obtained from liquid culture when bacteria and fungi grow together, and followed the expression level of HOG1, FUS3, and SGE1 genes of *P. viticola* considering a previous study carried out on *P. longicolla* which is in the same genus (Li et al., 2018) (Table 1).

**TABLE 1.**
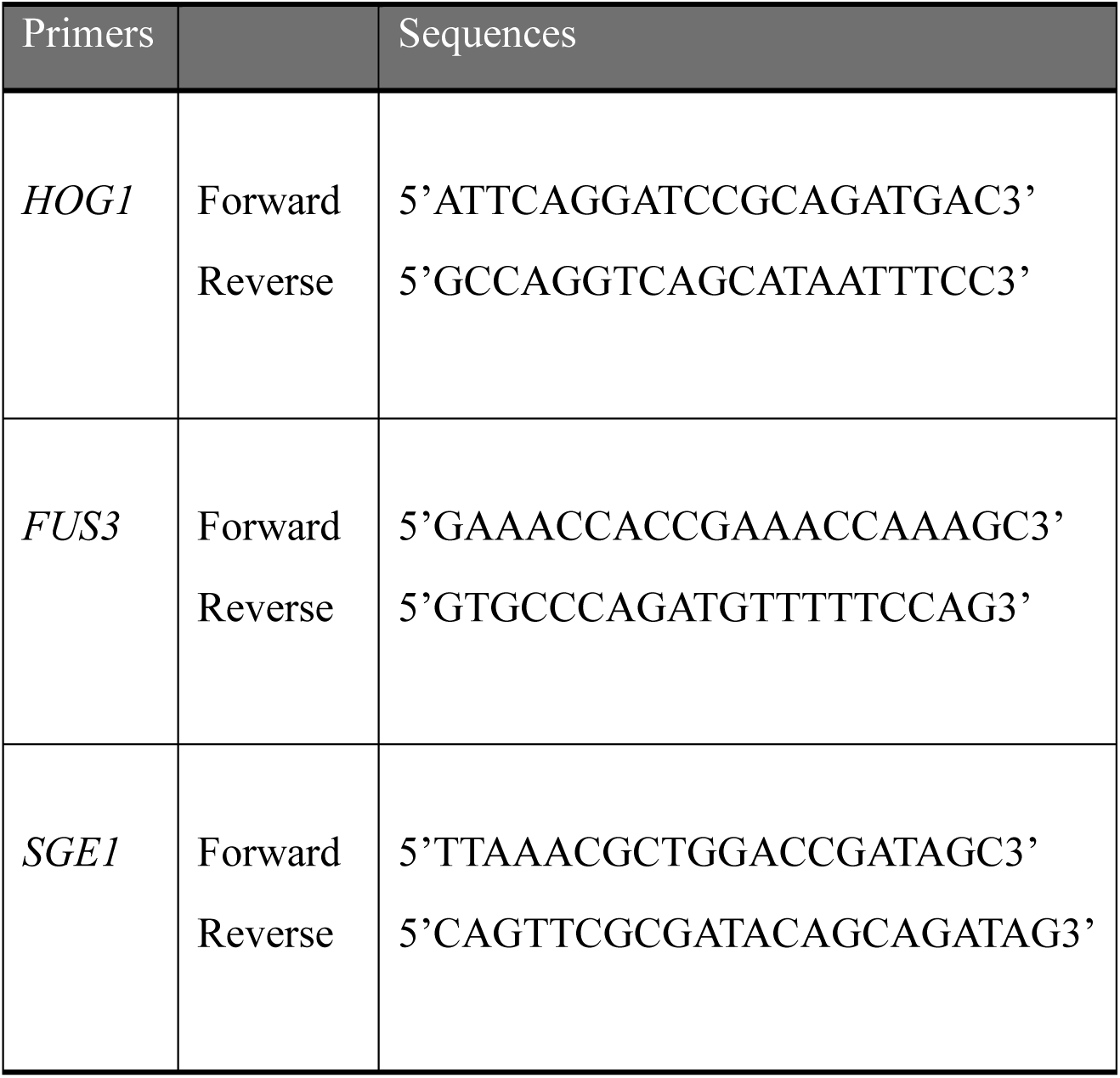
Primer sequences used for qPCR validation.

qRT-PCR analysis was performed on three major selected genes and validated as follows: The first cDNA strand was prepared using the iScript™ cDNA synthesis kit (Bio-Rad). qRT-PCR was performed using the SsoFast™ EvaGreen® supermix kit (Bio-Rad) in which each 20 μl of RT-PCR reaction mixture contained 25 ng of cDNA (equivalent of 25 ng RNA) and 0.125 μM of each primer. Three genes with non-altered expression (Actin) were selected as reference genes (AhActin-F-5’CTGAAAGATTCCGATGCCCTGA3’, AhActin-R-5’AACCACCACTCAAGACAATGTTACCA3’) (Du et al., 2022). Forward and reverse primer pairs (Table-1) for each gene were designed using Primer3 software (Rozen and Skaletsky, 2000) with an amplicon size of 90 - 200 bp (Supplementary Data S1). Real-time quantitative PCR was performed using the MyiQ™5 Real-Time PCR Detection System (Bio-Rad) using the following thermal cycle parameters: initial activation 100 at 95 °C for 30 s, followed by 40 cycles of denaturation at 95 °C for 5 s and annealing/extension at 58-60 °C for 10 s. Each run was followed by melt curve analysis. A standard curve was prepared for each gene using a cDNA sample. The relative expression of differentially altered genes was analysed using a free tool (https://goldbio.com/qpcr-and-rtqpcr-analysis-tool) according to Pfaffl (2001).

### 2.8 Molecular docking-based virtual screening for protein-ligand interactions

The experimental X-ray diffraction structure of HOG1 (high osmolarity glycerol 1), with its corresponding Protein Data Bank (PDB) identification number 1WFC, was obtained from the Research Collaboratory for Structural Bioinformatics (RCSB) PDB website (Wilson et al., 1996; Berman et al., 2003). The missing residues from the structure were added via PyMol’s builder plugin (open-source v2.5.0), loop regions where the residues were added, which were refined using MODELLER (v10.1) (Webb and Sali, 2016; Fiser et al., 2000; The PyMOL Molecular Graphics System, 2023). Subsequently, the structure was subjected to a purification process, in which all heteroatoms were removed, except for the atoms associated with the cofactors. In addition, polar hydrogen atoms were introduced where deemed essential, and Kollman charges were calculated (Morris et al., 2009). A grid box measuring 24 Å in the X-axis, 24.8 Å in the Y-axis, and 37.5 Å in the Z-axis was computed to encapsulate the active site of the 1WFC structure. Virtual screening was done utilizing the HOG1 structural model, evaluating its interaction with the thiamine ligand within the pre-determined grid box. Screening was conducted with an exhaustiveness level set at 64 using AutoDock Vina (v1.1.2) (Trott and Olson, 2010). The ligands exhibiting the highest affinity scores were subsequently selected using the same configuration.

### 2.9 Protein-ligand interaction profiling

The best dock pose of the ligand was loaded with HOG1 to PyMol, and all residues within 4 Å from the lead compound were visualized (i.e. all potential, hydrophobic interactions, hydrogen bonds, and ionic interactions) and evaluated. The manually predicted bonds were also cross-validated with the TU Dresden’s Protein-Ligand Interaction Profiler (PLIP) webserver, and only overlapping interactions were considered (Salentin et al., 2015).

### 2.10 Molecular dynamics analysis

Docking research often neglects the dynamic nature of proteins. For a comprehensive understanding of binding configurations and to assess system stability, it’s critical to perform molecular dynamics (MD) assessments. Such simulations offer computational approaches to explore the temporal dynamics of molecular arrangements (Ahamad et al., 2018, 2021). In our investigation, we utilized the Desmond component of Maestro’s academic edition to execute MD assessments (Schrödinger, 2021). This enabled us to scrutinize system interactions, structural alterations, and overall behavior between the ligand and protein. This investigation aimed to explore the stability and dynamic characteristics of a complex involving a protein and a ligand. We examined the protein PDB ID 1WFC along with thiamine as the ligand under simulation conditions. Our objective was to understand how ligands impact protein stability and to identify the protein sectors that exhibited the most significant variations. Our simulations were conducted in an NPT ensemble at 300 K for 100 ns. The OPLS-AA force field was employed for the complex, and ligand parameters were generated using Maestro’s LigPrep module (Schrödinger, 2021). The NPT ensemble is a form of MD simulation that accounts for the number of particles, pressure, and temperature within the system. This allows for a constant volume and temperature setting where the pressure can vary, making it useful for studying systems in a solution-based and more lifelike simulated setting. During the simulation, we kept track of the root mean square deviation (RMSD) for both protein and ligand. We also measured the protein’s RMSF to detect localized alterations along its chain. The RMSF metrics for individual residues were observed throughout the simulation, allowing us to identify the most variable regions within the protein structure. In addition, we mapped out contacts between the ligand and protein to identify the specific residues involved in ligand binding, marking these areas with green vertical lines for easier visualization. We also continuously observed the protein’s secondary structure, which refers to local folding patterns such as alpha-helices and beta-sheets. These structures are critical for protein functionality, and their alterations can influence its activity. Finally, we documented the existence of counterions and the concentration of salts in the solvent, as these can significantly affect the system’s dynamics. To summarize our MD simulation offered valuable insights into the stability and dynamics of a protein-ligand complex under specific conditions, including the monitoring of RMSD and RMSF values, the identification of residues interacting with the ligand, observations of the protein’s secondary structure, and the recording of counterions and salt concentrations in the solvent.

## 3 Results and Discussion

### 3.1 Identification of MM19 and Antagonistic assay using single layer assay

The Biolog GEN III results have shown that the bacterial strain belongs to *Bacillus safensis* genus (Supplementary Figure S1). Further analysis was carried out considering these results (Supplementary Table S1). A significant decrease in the radial expansion of *P. viticola* became clear upon the introduction of *B. safensis* MM19 from day 7 onward, and this inhibitory effect endured throughout the entire experimental duration (Supplementary Figure S2, Supplementary Table S2).

### 3.2 Transcriptome profiling of *B. safensis* after *P. viticola* interaction

The obtained RNA-seq data were deposited in the DDBJ Sequence Read Archive at the DNA Data Bank of Japan (http://trace.ddbj.nig.ac.jp/dra) under accession numbers SAMD00731600 (BioProject; PRJDB17356). The analysis generated 1.4 million raw reads for each sample, and 80.0% of them were correctly mapped to the *B. safensis* reference genome of NCBI, which contained 3996 annotated genes. The mean CPM values and k-values in the samples are shown in Supplementary Data S2. In this study, after *P. viticola* interaction, the genes showed 2-fold increase in upregulation and downregulation (Table 1) (Supplementary Figure S3). From 1004 genes, 527 genes were upregulated and 477 were downregulated (Figure 3, Figure 4 and Supplementary Data S2). Totally, RNA-seq analysis indicated 3398 genes of bacteria involved in this interaction with fungi when they growth together in liquid culture.

**FIGURE 3.**
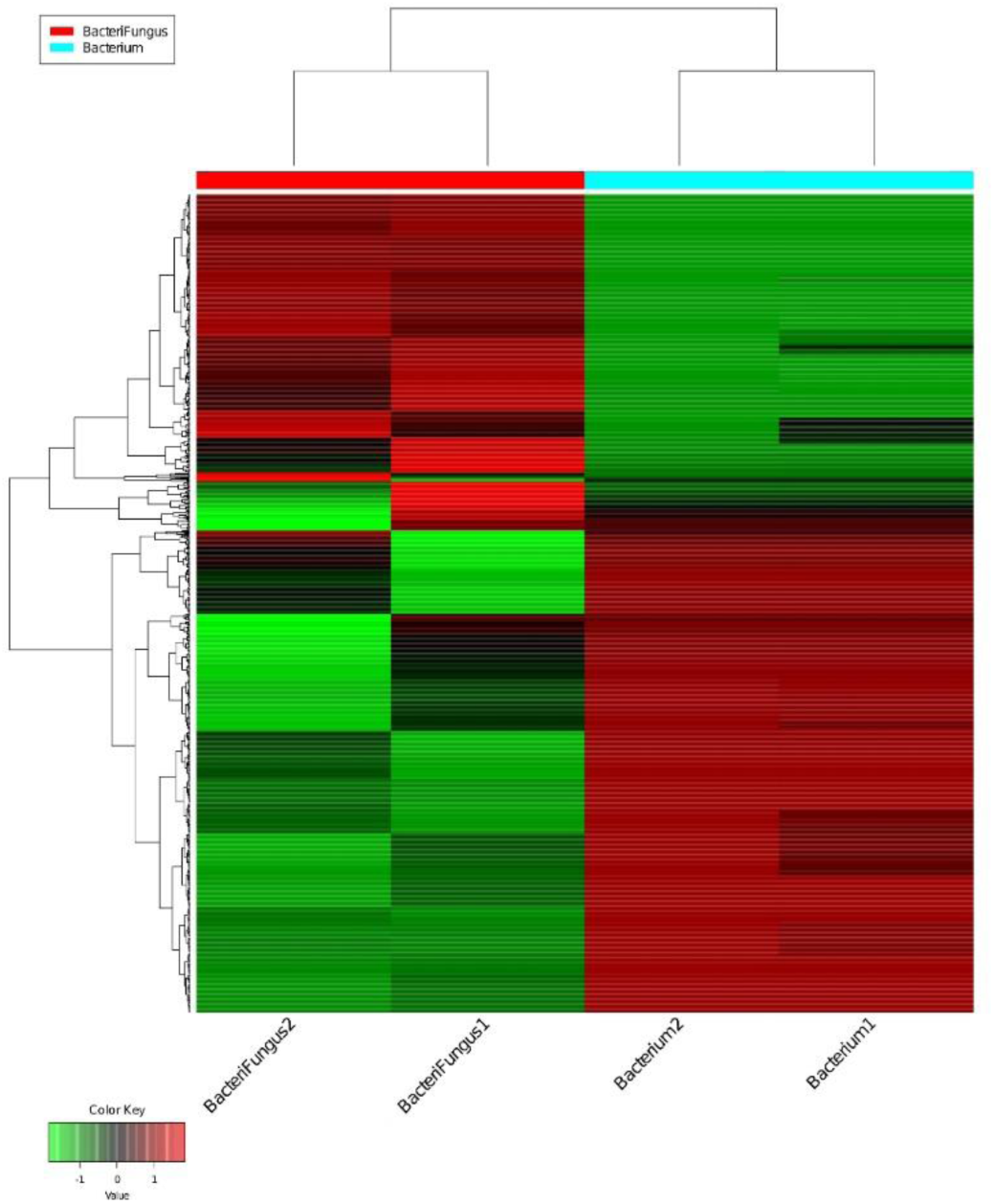
Transcriptomic data obtained from the flasks of bacteria-fungi interaction groups and bacteria alone. Transcriptional levels of the relevant genes in *Bacillus safensis* MM19 are shown as a heat map, based on the ranking of expression intensity in the whole genome. The color of each square represents the strength of gene transcription.

**FIGURE 4.**
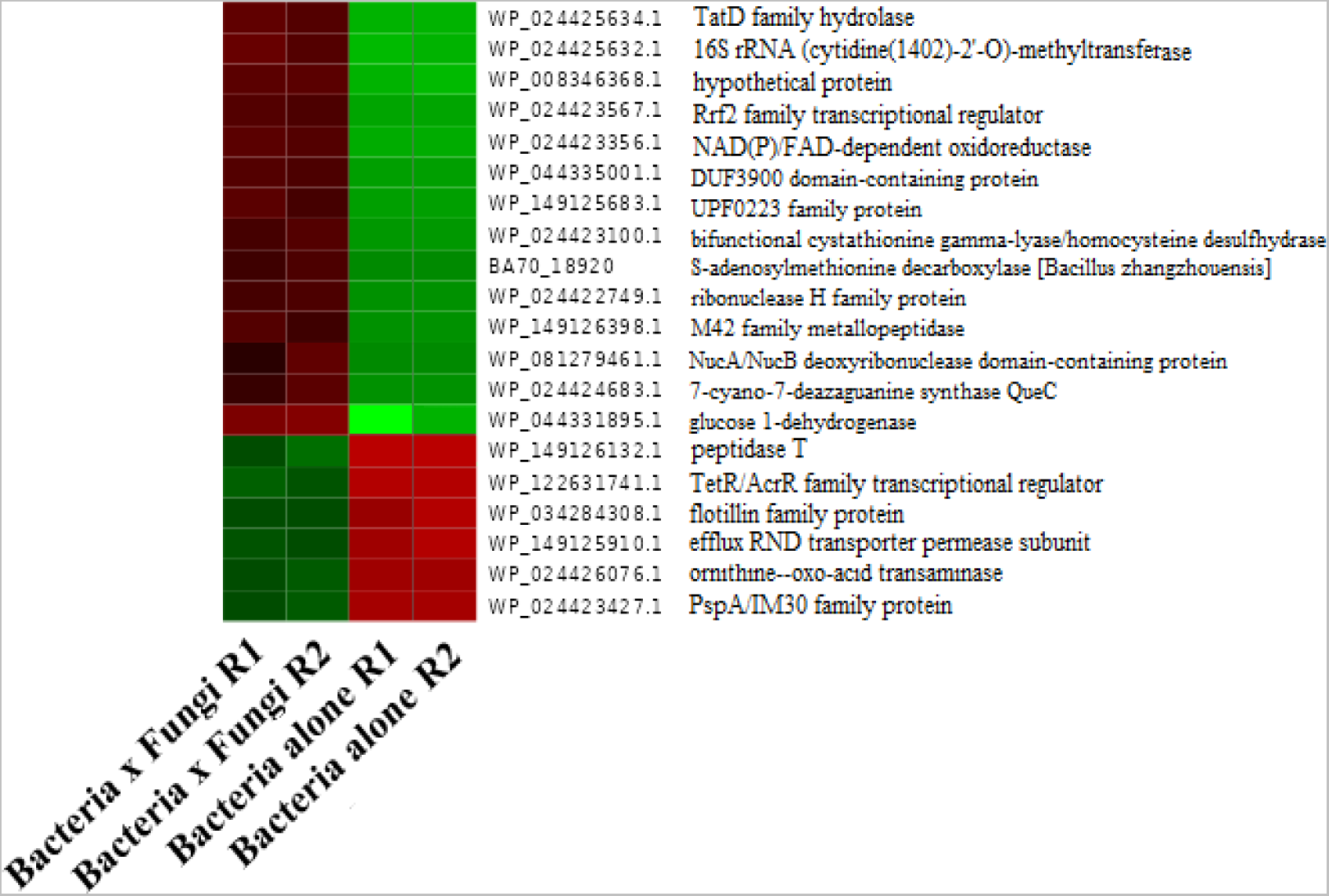
The most variable top 20 genes in bacterial transcription during the bacteria-fungi interaction according to logFC values.

The transcriptomic data obtained from bacteria show changes up- and downregulation. The expression profile has changed depending on stress conditions in bacteria and varies in RNA samples obtained from bacteria-fungi interactions and bacteria alone groups. The expressed genes demonstrated differentiation. The genes upregulated in bacteria growth in liquid culture medium showed downregulation when the bacteria growth with *P. viticola*, and vice versa, the genes that were downregulated before showed upregulation ascending when the bacteria growth with *P. viticola* (Figure 3, 4). The S-adenosylmethionine decarboxylase genes showed high upregulation (Figure 4). In a previous study, proteomics analysis also indicated up regulation of S-adenosylmethionine (SAM)-dependent methyltransferases in a bacterial strain belonging to *Bacillus* species (EU07) (later identified as *Bacillus amyloliquefaciens*) when it encountered *Fusarium oxysporum* radicis *lycopersici* (Baysal et al., 2008). Correspondingly, in the present study, RNA expression analysis indicated a decrease of S-adenosylmethionine decarboxylase converting 1-aminocyclopropane-1-carboxylate (ACC) synthase into S-adenosylmethionine (AdoMet) when MM19 encountered *P. viticola* (Figure 4). This result could be linked with the substitution of S-adenosylmethionine with SAM-dependent methyltransferases, which could allow bacteria to utilize this pathway as an alternative response to *P. viticola*, which could be linked with quorum sensing and common virulence behavior of the bacteria to target pathogen. It is also noteworthy that these genes could also be involved in riboswitches, which are highly conserved metabolite-sensing noncoding RNAs responsible for the regulation of gene expression. They are mostly found in the untranslated regions of mRNAs and can bind to small-molecule metabolites directly, thereby regulating the expression of metabolic-related genes (Coppins et al., 2007). Riboswitches can sense various cellular metabolites whose gene products are always encoding with the riboswitch downstream sequence. These metabolites include coenzymes, amino acids, metal ions, and nucleobases (Roth and Breaker, 2009). Thiamin diphosphate riboswitch is one of the most widespread riboswitches. Members of this class regulate the expression of genes involved in the biosynthesis, salvage and transport of thiamin and its precursors (Kubodera et al., 2003; Kazanov et al., 2007). Studies by Schyns et al. (2005) on *Bacillus subtilis* confirmed that the mechanism of thiamine gene regulation is controlled via riboswitch. RNA analysis results showed that ribosomal protein translation significantly increased and also energy consumption increased with ATP synthesis (Figure 5).

**FIGURE 5.**
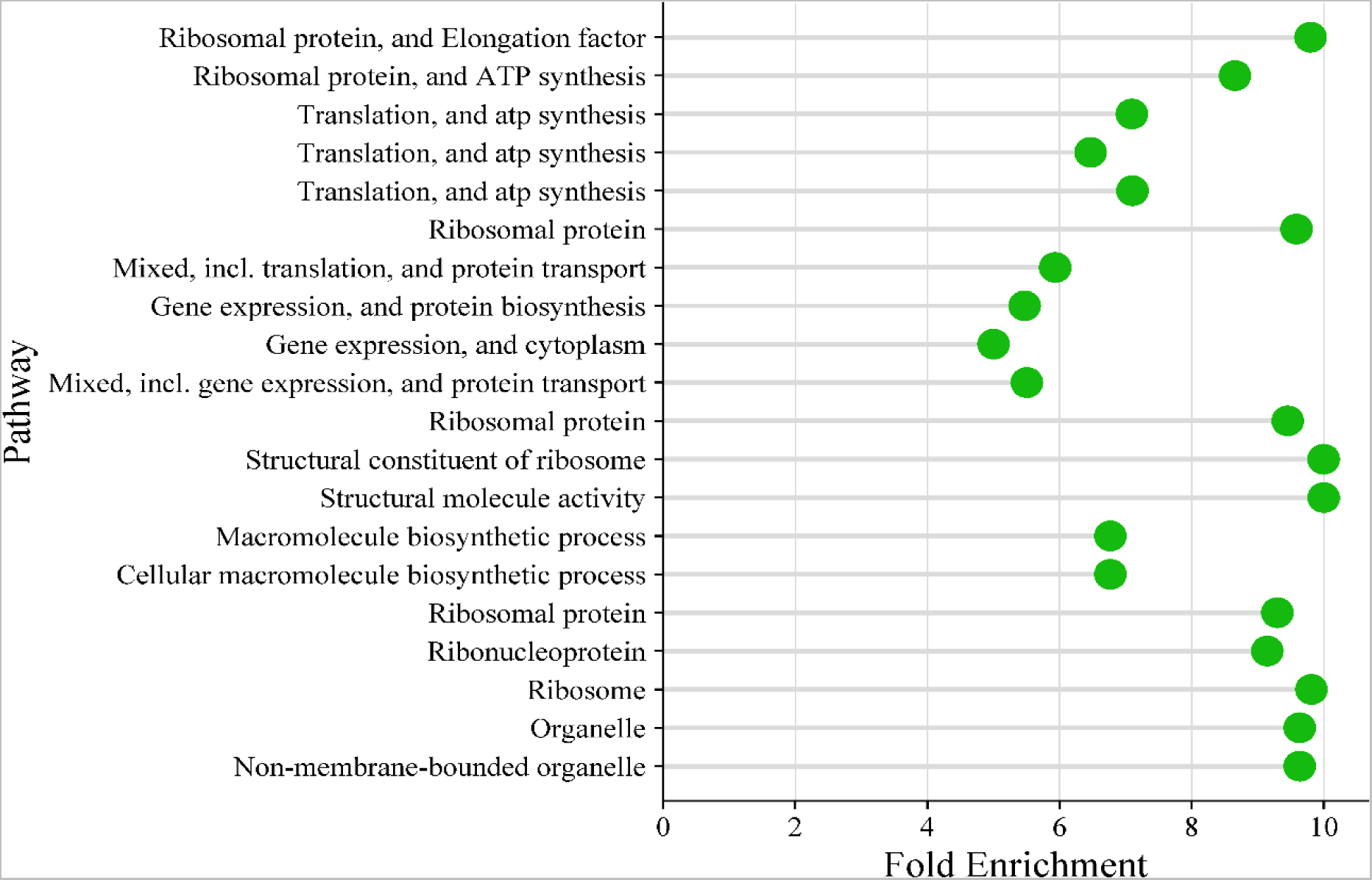
The chart shows hierarchical clustering tree summarizing the correlation among significant pathways listed in the enrichment table. Pathways with many shared genes are clustered together. Bigger dots indicate more significant P-values.

The prominent genes, ranked according to their degree of importance in the interaction between bacteria and fungi, are listed in the Table 2, separated by functional groups. These results show which genes of the bacterium that encounters the fungus have a respond at transcriptional level under biotic stress. Findings regarding cellular metabolism showed that the response of MM19 to *P. viticola* was increased the gene expression level with many metabolic bioprocesses and specifically resulted in enhanced protein synthesis (Figure 6). We observed that particularly, the ribosomal protein and transcriptional process show ascending depending on biotic stress on bacteria. Considering the genes given in Table 2, common expressions occurring in the genes of bacteria can be observed that based on this finding, we could use these data as a parametric measurement for qPCR-based methods to understand of antagonistic behavior involving responsible genes for further studies.

**FIGURE 6.**
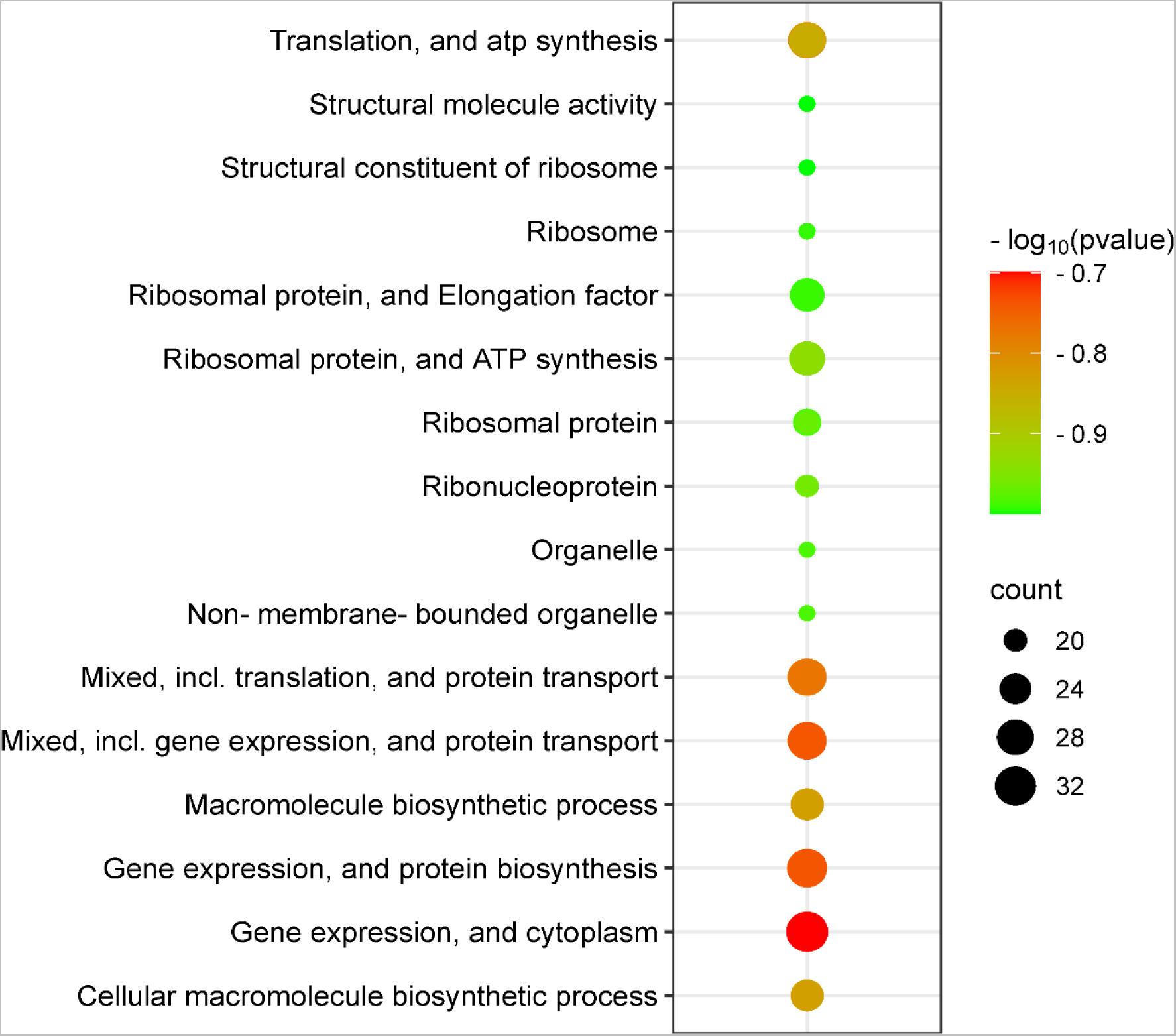
The illustration shows gene enrichment analysis related to cell metabolism pathways depending on RNA-seq analysis counts that reveals up-regulated or down-regulated genes during bacteria and fungi interaction.

**TABLE 2.**
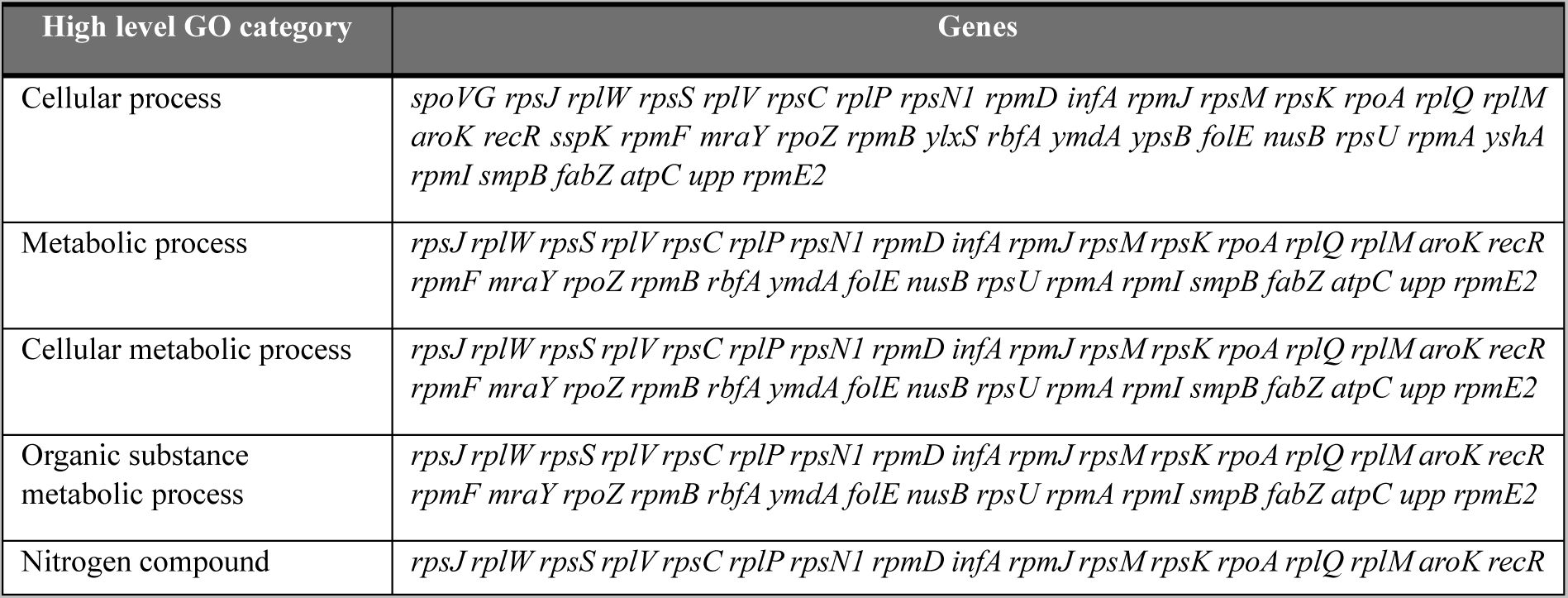
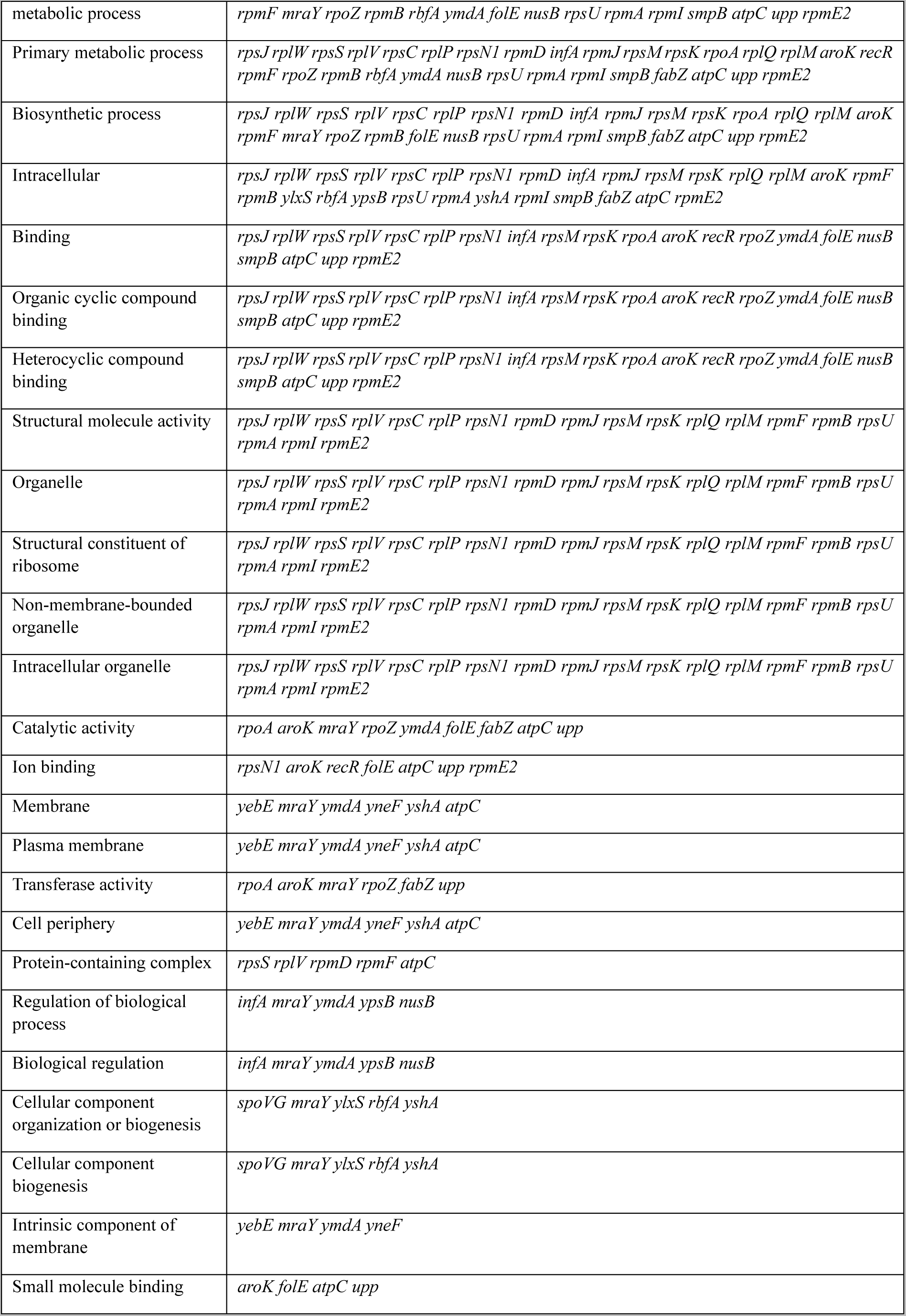
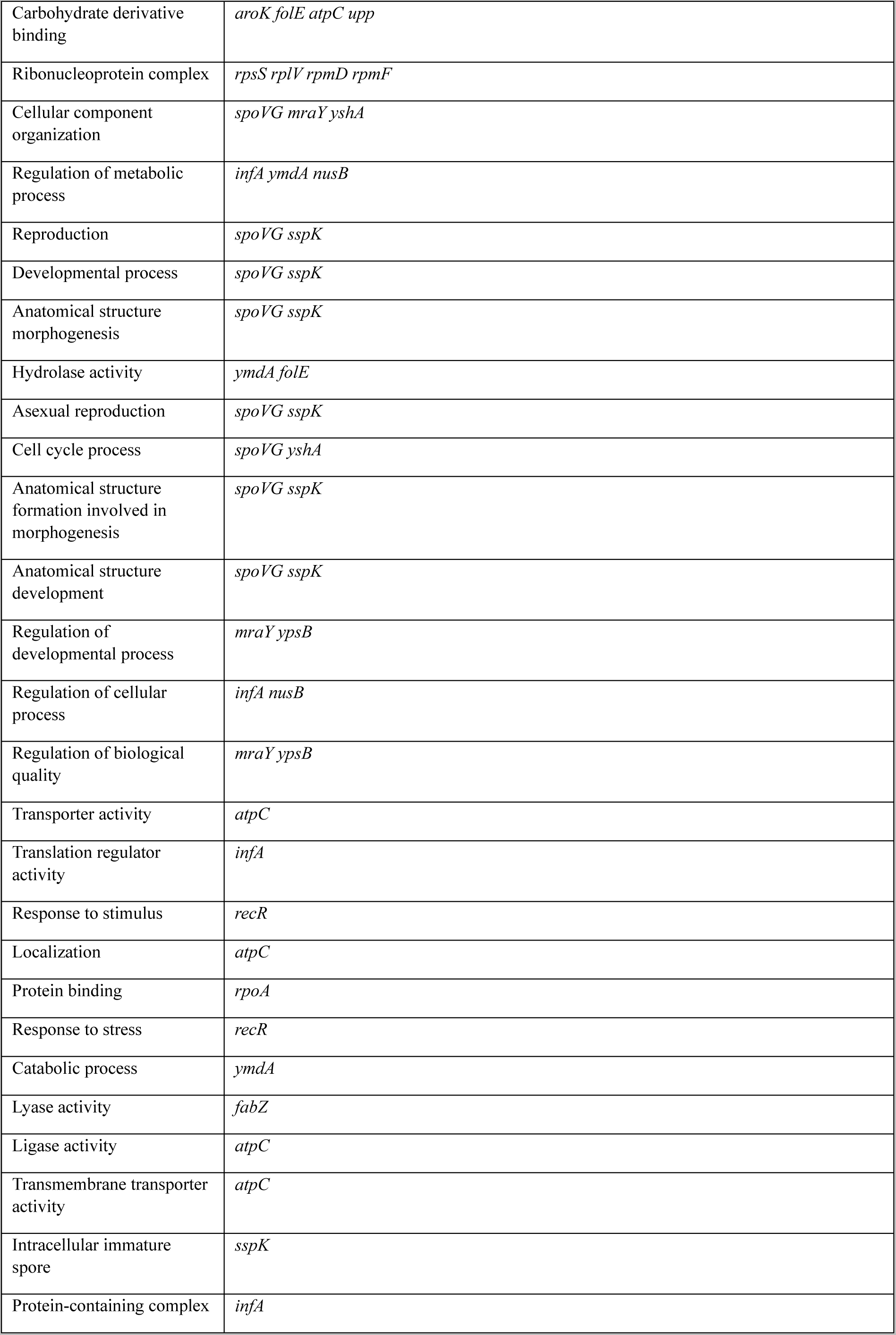
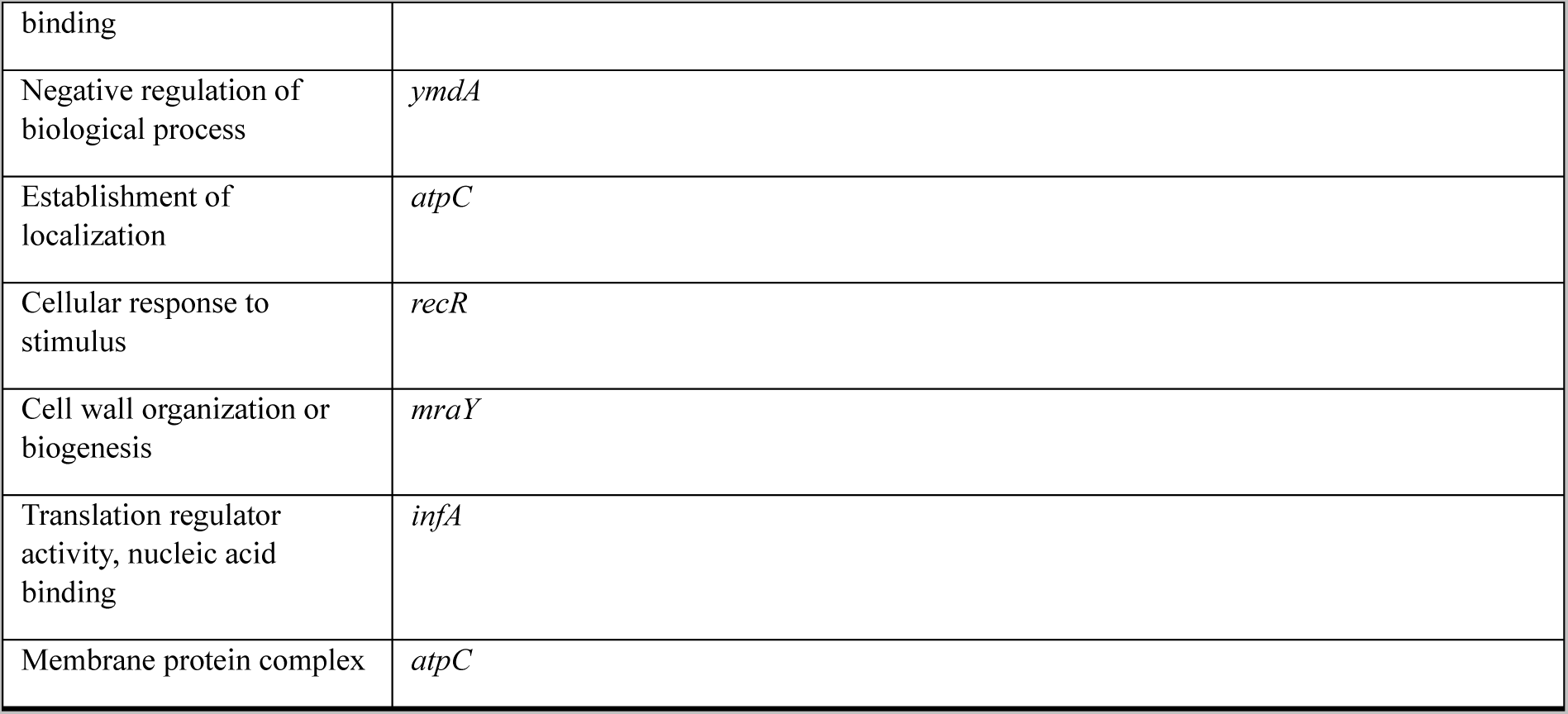
Genes are grouped by functional categories defined by high-level GO terms according to RNAseq output showing up-down regulation.

There is no genomic database for the comparison of the expressed specific genes in *B. safensis* for gene enrichment analysis in ShinyGO, therefore we have used *B. subtilis* strain 168 as a reference. The p values of the biosynthesis and metabolic events that stand out in the enrichment analysis have been listed in the appendix, as visualized in ShinyGO (Supplementary Data S3, Supplementary Data S4). Since the bacterium varies the regulation of several genes depending on their functions, it is more valuable to interpret the clustering of these genes. The gene enrichment analysis reveals the major metabolic pathways of up-regulated or down-regulated genes during bacteria and fungi interaction (Figure 7). In this analysis, the oxidoreductase activity acting on CH-NH^2^ group of donors, oxygen as receptor, FAD dependent oxidoreductase and FAD binding processes play a major role of total metabolic processes. Also, most of the genes involved in thiamine metabolic pathways were highly expressed during the interaction with fungi (Figure 7).

**FIGURE 7.**
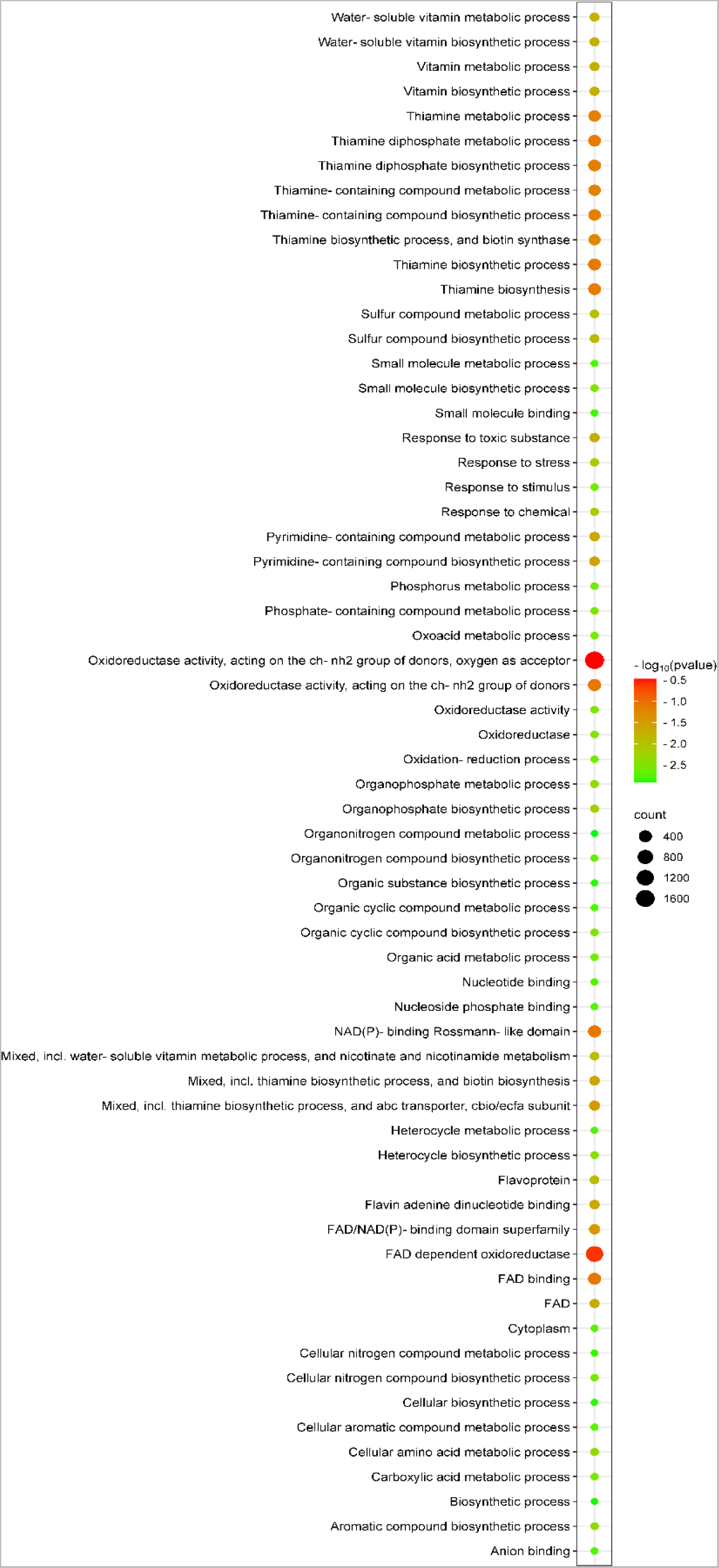
The illustration shows gene enrichment analysis related to metabolic pathways depending on RNA-seq analysis counts that reveals up-regulated or down-regulated genes during bacteria and fungi interaction.

Specifically, oxidoreductase consists of several different enzymes such as peroxidase, reductase, dehydrogenase, oxidase, oxygenase, and hydroxylase. It can take part in both anaerobic and aerobic metabolisms (Younus, 2019). Oxidoreductases catalyze oxidation and reduction reactions that occur within the cell. However, it often needs cofactors such as nicotinamide adenine dinucleotides (e.g., NAD^+^/NADH) and flavines (e.g., FAD/FADH^2^) in the reactions. In fact, nicotinamide adenine dinucleotides are required by about 80% of oxidoreductases. Several NAD (H) regeneration systems have been developed; the most widely used is formate-formate dehydrogenase (FDH) system (Paul et al., 2019). In the sight of these knowledge, our RNA-seq data indicated when bacteria encountered with fungi, it increased oxidation metabolism than before.

Diaporthe (*Phomopsis*) fungi generate natural products such as pyrones, polyketides, alkaloids, and terpenoids. Most of these natural products show antibacterial, anti-inflammatory, and/or cytotoxic activity (Jiang et al., 2023). One of these products is Trichothecene which is one of the major classes of mycotoxins, has a common 12, 13-epoxytrichothec-9-ene structure, in which the epoxy group is the active site leading to oxidative radical formation (McCormick et al., 2011; Mostrom and Raisbeck, 2007). The oxidoreductase dependent metabolic pathways could be over expressed to overcome the oxidative radicals secreted by fungi (Figure 7). In a previous study, Afsharmanesh et al. (2018) found oxidoreductase from *B. subtilis* UTB1 which is able to degrade aflatoxins, very toxic and hazardous secondary metabolites produced by a specific strain of *Aspergillus* species, e.g., *Aspergillus flavus* and *Aspergillus parasiticus*. Their study successfully degrades aflatoxin by various oxidoreductase activity and reduces the growth of fungal biomass. Therefore, we can assume bacterial oxidoreductases of MM19 could have a major role for degradation of the toxic metabolites of the fungus by *Bacillus* species.

Thiamine (vitamin B1) is an essential cofactor for carbohydrate metabolism (Hwang et al., 2017; Chen et al., 2019) involving biosynthesis and degradation of the branched-chain amino acids (Schauer et al., 2009). Therefore, the availability of thiamine has a key effect on the growth of many organisms, e.g., *Xanthomonas oryzae* pv. *oryzae* (Yu et al., 2015) and *Listeria monocytogenes* (Madeo et al., 2012). Noteworthy, the studies of Liu et al. (2022) indicated thiamine is essential for virulence and survival of *Pseudomonas syringae* pv. *tomato* that we can assume the transcriptional activation of thiamine related genes in MM19 has a major role of challenge during the *P. viticola* interaction. Also, there is no data on the antifungal effects of the thiamine derivatives. From this point of view, we have investigated the docking simulation of a virulence protein of *P. viticola*, HOG1 and thiamine interaction.

### 3.3 Enrichment gene interaction map and network analysis

Figure 8 shows metabolic map of the pathways responsible for thiamine biosynthesis. The metabolic network was constructed based on KEGG pathway analysis using Cytoscape (Supplementary Data S5; S6; S7; S8, Supplementary Figure S4; S5; S6). Four modules were partitioned according to their functions, respectively. The illustration shows different metabolic modules: 1, Purine pathway, 2, Cysteine and Methionine pathway, 3, Glycolysis/Gluconeogenesis pathway module, 4, Phenyalanine, Glycine pathway.

**FIGURE 8.**
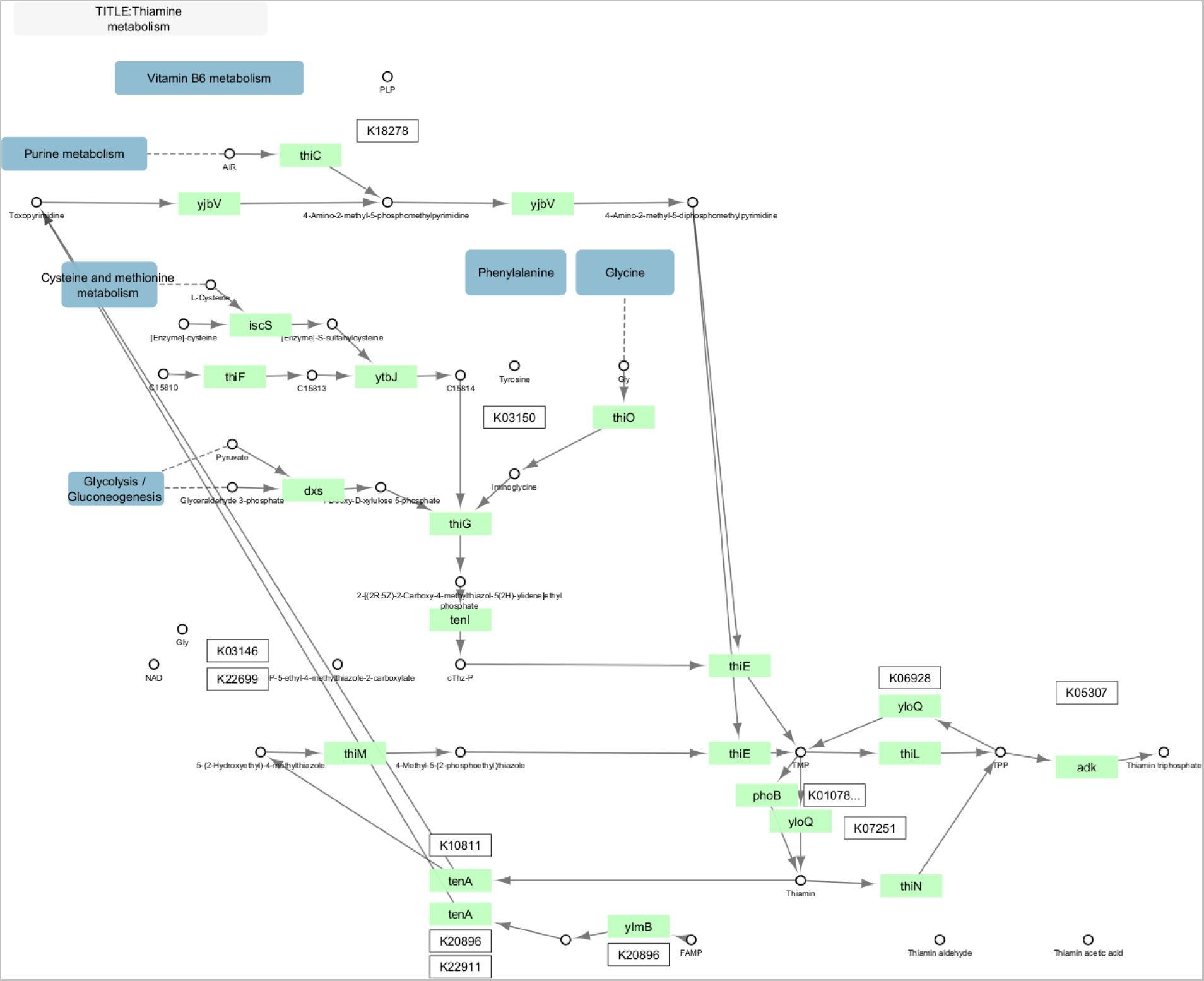
The illustration shows expression analysis of network using Cytoscape. The network was constructed considering major variable genes obtained with RNA-seq analysis.

### 3.4 qRT-PCR analysis on selected 3 major genes responsible for the virulence

Even we tried obtaining RNA sample of *P. viticola* through liquid dual culture, we were not able to supply sufficient uncontaminated RNA quantity without bacterial DNA contamination, which was necessary for library construction of *P. viticola*. Therefore, we could not obtain raw RNA-seq data from fungi as we have requested. Instead of RNA-seq, we selected 3 major genes playing role in virulence of *Phomopsis longicolla* which has the most genetically close and available corresponding genes of *P. viticola* (Li et al., 2018).

In the qPCR experiments conducted with 4 replicates, the higher expression of HOG1 than the others showed that the interaction of HOG1 with thiamine, have much more considerable case that should be focused on terms of pathogenic growth inhibition by antagonistic bacteria (Figure 9) (Supplementary Data S9).

**FIGURE 9.**
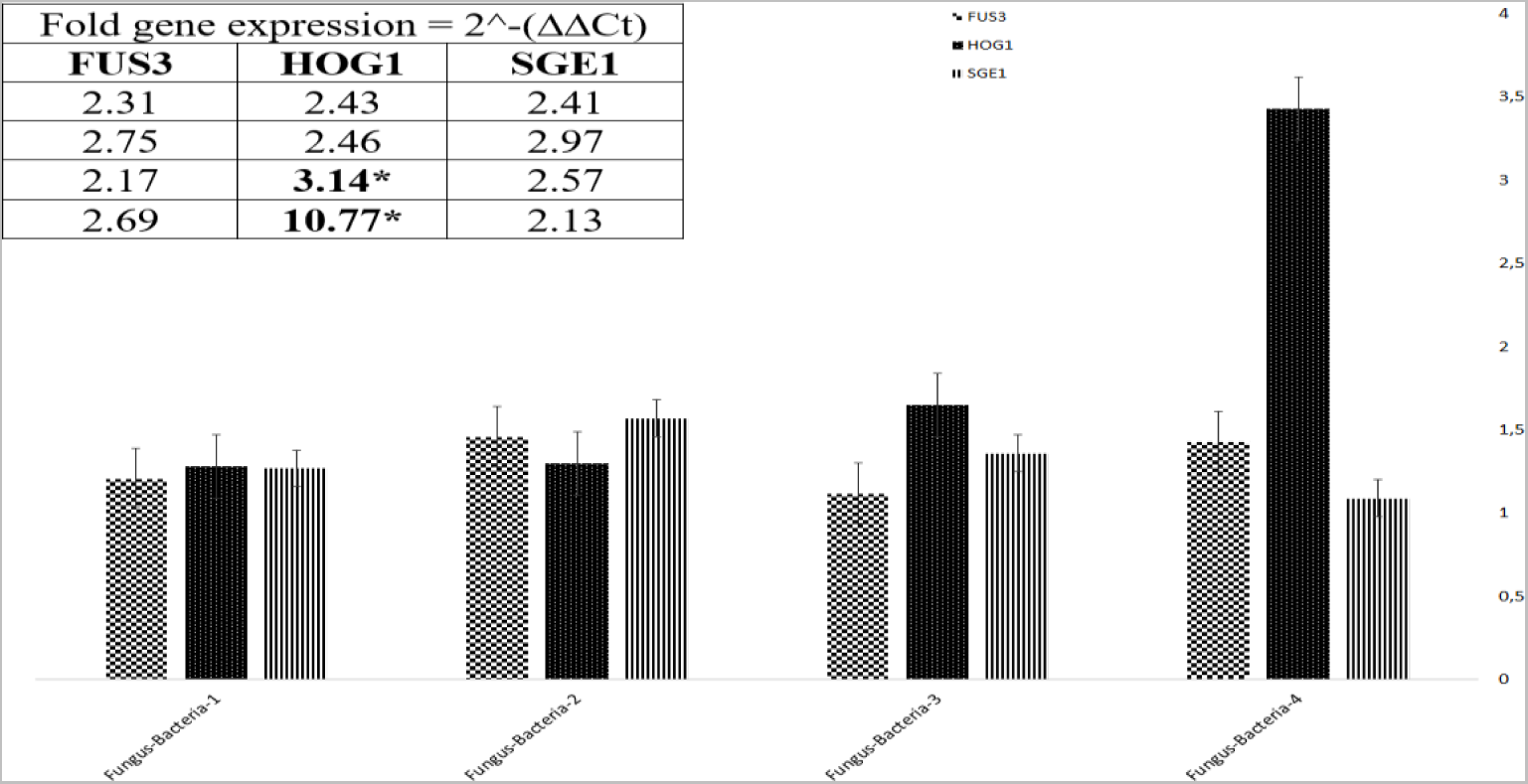
qPCR validation of selected differentially expressed genes with four replicates. The fold gene expression has given as 2^-(ΔΔCt) value compared to housekeeping gene ß-Actin.

### 3.4 Molecular docking-based virtual screening for protein-ligand interactions

The process of virtual screening assessed the ligand’s (thiamine) affinity to the HOG1, with a specific criterion set at a binding energy threshold ≥ -6.5 kcal/mol. This stringent threshold is indicative of substantial binding strength and the potential for effective ligand-mediated inhibition of the target protein.

### 3.5 Protein-ligand interaction profiling

The interaction between the HOG1 catalytic site with the divalent cation and the thiamine (as shown in Figure 10, Supplementary Data S10) was closely analyzed and visualized. Figure 10A illustrates the comprehensive depiction of potential hydrophobic interactions and hydrogen bonds established between individual residues and the lead compound within the active site of HOG1. Additionally, the interacting residues originating from the thiamine backbone are meticulously expanded to enhance the visualization of the interacting atoms.

**FIGURE 10.**
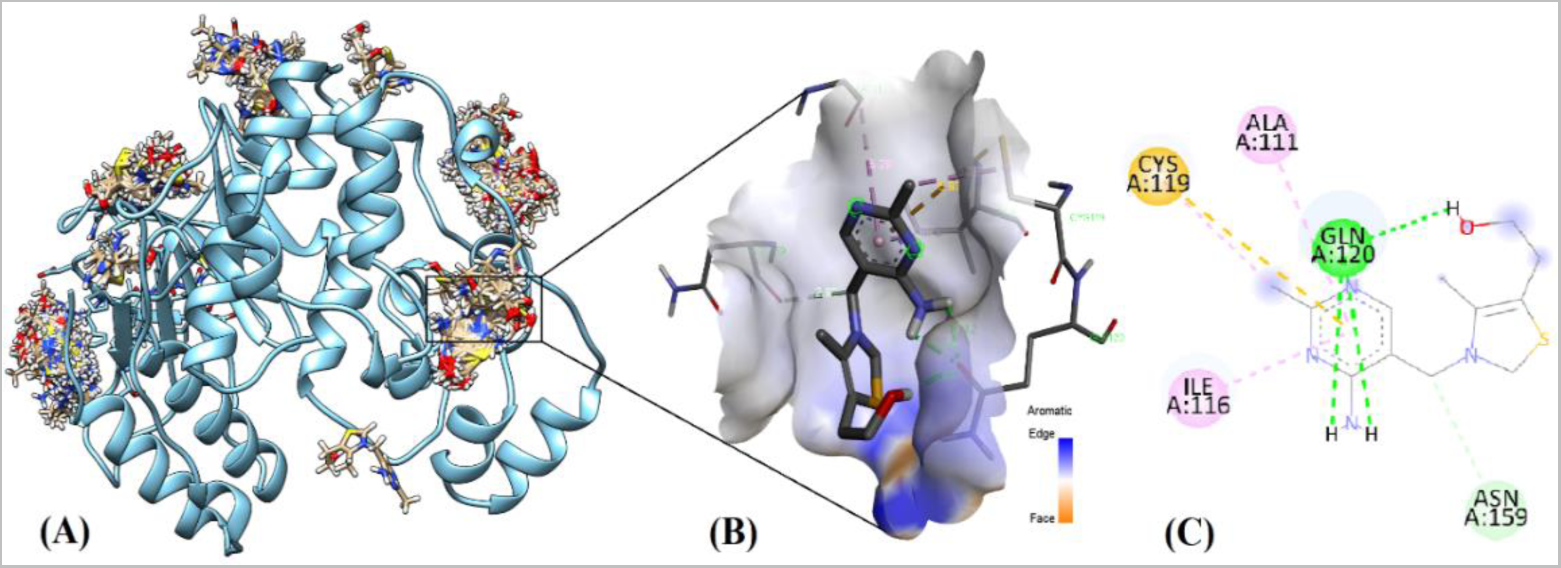
(A) The complete docking conformation of thiamine (depicted in blue and red) with the transmembrane domain of HOG1, characterized by helical structures, was investigated, focusing multiple thiamine compound binding events on the target protein. The highest binding energy between HOG1 (receptor) and thiamine (ligand) and its coordinates on receptor shown in **(B)** 3D and **(C)** 2D.

### 3.6 Molecular dynamics analysis

The stability of the interaction between HOG1 and thiamine was systematically investigated by simulation of 100 ns molecular dynamics in Figure 11. The RMSD values were employed to assess the equilibrium attainment within the simulation and to discern any notable conformational alterations. Residues 50-60 of the protein are the most fluctuating regions and are effective in ligand binding. The simulation did not exhibit significant alterations in the secondary structure of the protein over its simulation. The protein–ligand RMSD value showed that the protein and ligand were stable throughout the simulation. RMSD value varied between 3.2 and 5.6. The RMSD value of the ligand indicates that the ligand fluctuates more throughout the simulation. The RMSD value ranges between 1 Å and 4 Å. The protein jumped around 50 ns. This finding suggests that the ligand has switched to an alternative binding mode. Therefore, the stability of the ligand could have decreased at this point. In conclusion, our MD results show that the protein and ligand remain stable throughout the simulation, but the ligand switches to different binding modes at some points. This could affect the binding stability of the ligand. In the ligand RMSF graph, ligand atoms fluctuate in different ways (Supplementary Data S11). Atom number 17 has the highest RMSF value. This atom could play an essential role in the interaction of the ligand with the protein. Additionally, the RMSF value of the ligand varied between 1 and 4 Å (Figure 12A). The most robust interaction between protein and ligand was with the ASP residue. This relic has a percentage value of 100%. This means that this residue is in constant contact with the ligand. This residue could play an essential role in binding the ligand to the active site of the protein. In Figure 12B (contacts), there are also weaker interactions between the protein and the ligand. These are the ones with percentage values lower than 50%. These interactions mean that the ligand temporarily comes into contact with different regions of the protein and can affect the binding stability of the ligand (Supplementary Data S11).

**FIGURE 11.**
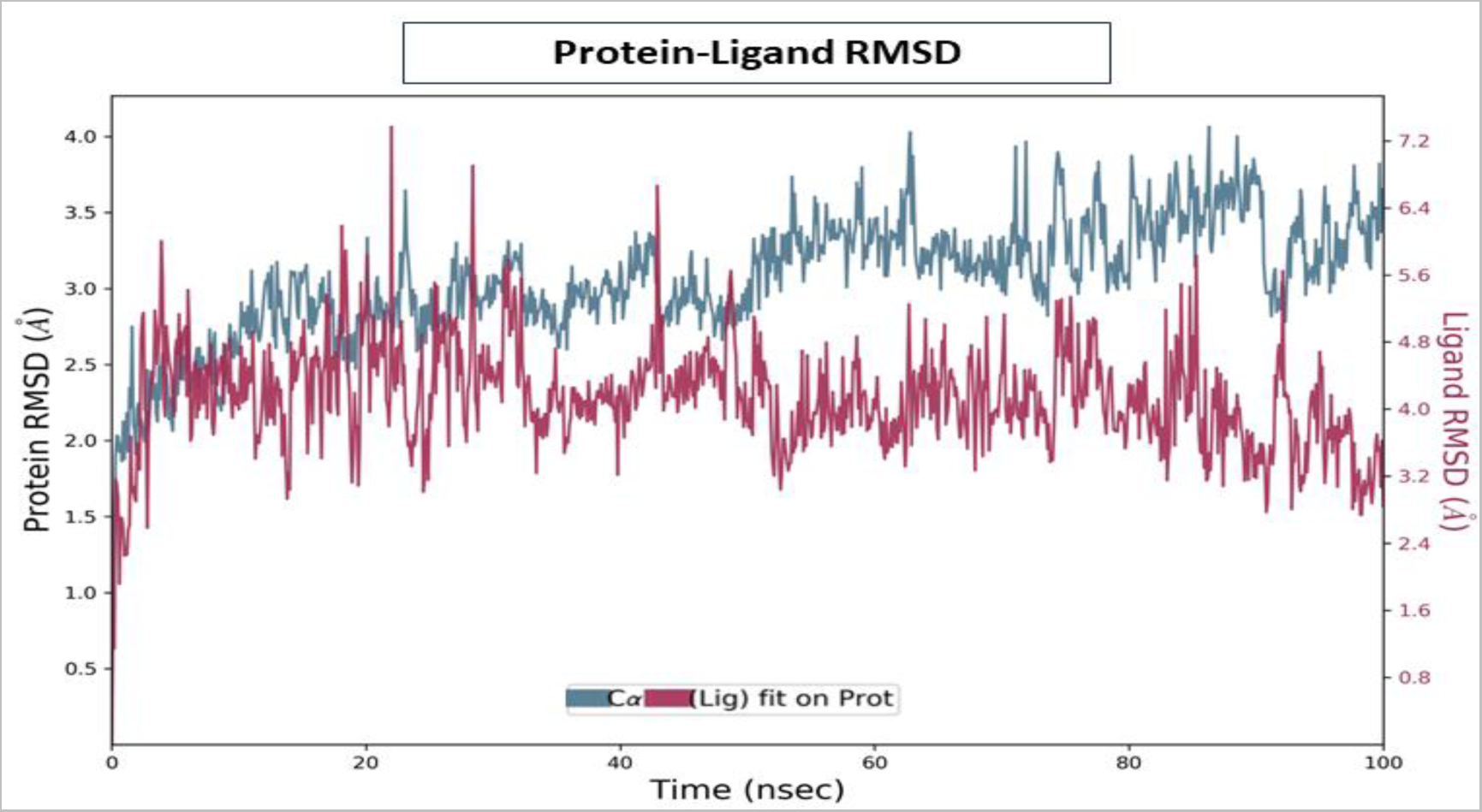
HOG1 and thiamine simulations were conducted with reference to the initial frame throughout the entire 100 nanosecond production simulation. The cyan line in the graph depicts the trajectory of HOG1 motion. Notably, reduced variability along the Y-axis indicates a lesser degree of deviation from its initial docked position. The red line in the plot represents the root mean square deviation (RMS) of the thiamine backbone, illustrating the displacement of HOG1 during the course of the simulation.

**FIGURE 12.**
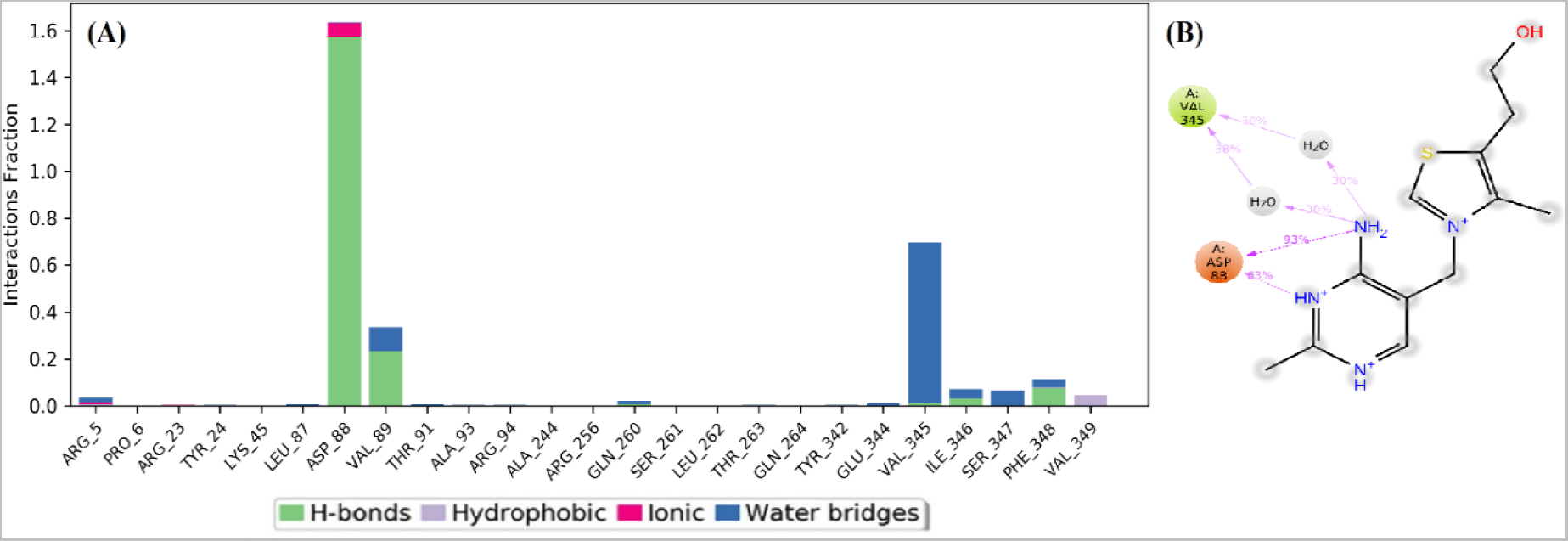
(A) The histogram illustrating the occurrences of protein-ligand contact along the molecular trajectory is presented. (B) The structural configuration of the protein-ligand complex system was derived through a molecular dynamics simulation, specifically focusing on the two-dimensional interaction mode. Notably, hydrogen bonds, salt bridges, and π-cation interactions are respectively delineated in the visual representation using green, pink, and dark green lines.

HOG1 is a mitogen activated protein kinase and has major roles in fungal cell growth. Mitogen-activated protein kinases are eukaryotic serine-threonine kinases that mediate vital intracellular programs like proliferation, differentiation, survival, and apoptosis (Morrison, 2012). Also, the studies on *Saccharomyces cerevisiae* by Brewster and Gustin (2014) revealed HOG1 as a central signaling mediator during osmoregulation. Due to signaling specificity properties of the kinases involving response and substrate specifity behavior, HOG1 and FUS3 have attributed as regulator proteins responsible for protein-protein interaction and cell recognition (Mody et al., 2009). In the sight of our findings, it can be assumed that bacteria have targeted these two specific kinases by emitted effectors to diminish the pathogenic behavior of *P. viticola*. The bacteria’s thiamine production could be linked to HOG1 which resulting in impairment on mitogen kinase activated protein responsible for the signal communication of the cell wall. The targeting could be the antagonistic strategy of the bacteria to distract of the pathogen for diminishing its signal transduction causing specific mRNA transcription leading to biosynthesis of metabolites. Hence, as described by Siemieniuk et al. (2016) thiamine antivitamins could be considered as antifungal agents which suggests such new thiamine derivatives targeting HOG1 can also be employed and tested against *P. viticola* as a pathogen growth inhibitor compound.

Correspondently, independent studies revealed that thiamine triphosphate can act as a signaling molecule in adaptation of bacteria to stress conditions (Bettendorff and Wins, 2013). Adenosine thiamine triphosphate has been suggested to have a role in response to abiotic stress (Gigliobianco et al., 2010). Moreover, adenosine thiamine triphosphate is known to be a regulator for the activity of membrane adenosine thiamine triphosphate transporter (Nabokina et al., 2014) and poly(ADP-ribose) polymerase-1 (Tanaka et al., 2011). Given MM19 expressed higher levels of these genes responsible for thiamine biosynthesis (Figure 7), the prominent property of the bacteria could be linked with its inhibitory effect against pathogenic fungi.

Yu and colleagues (2019) recently used genome-wide transcriptome profiles to study how *Bacillus subtilis* C232 lipopeptides inhibit microsclerotia formation in *Verticillium dahliae*. The lipopeptides inhibited expression of genes associated with secondary metabolism, protein catabolism, and the high-osmolarity glycerol response signaling pathway. The suggested data provided insights into the activation of genes responsible for the assimilation of fungal-derived compounds and the potential synthesis of an unverified antifungal agent. Also, *B. safensis* MM19 could produce various genes encoding protease and a gene encoding esterase/lipase (Supplementary Data S3). These bacterial enzymes could inhibit the cell wall of the pathogen on account of antagonistic behavior. Since the cell membrane (plasma membrane, plasmalemma) of *P. viticola* involved essentially a lipo-protein structure, with a molecular organization that coincides with the principles of the fluid mosaic model (Coppin et al., 1997), we can assume these bacterial enzymes and especially lipopeptides also play role in antagonistic mode of action.

Bacterial genes encoding products of similar adaptational functions are frequently coregulated (Figure 6, 7). This organization ensures the balanced production of all proteins necessary for adaptation in terms of the antagonistic competition. Reorganization of the expression profile depending on signal transduction could be a featured case for the competition between secreted metabolites by antagonist bacteria and the stress factors affecting the pathogen, which limits its survival (Seshasayee et al., 2006). The further studies on integrated network model involving cellular behavior will enhance our profoundly understanding and knowledge on the correlation between genetic variations and metabolic pathways (Goelzer et al., 2008). The RNA-seq analysis is a key approach to determine the genes responsible for the response of bacteria under various stress conditions resulted from abiotic and biotic sources. This technique presented in our study could pave for unveiling microbial behavior of each strain with antagonistic potential when it encounters with target pathogen, which could be used to study in further investigations.

## 4 Conclusion

This study investigated whole gene expression profile in *B. safensis* MM19 during interaction with *P. viticola* in a liquid medium through RNA-seq analysis. The simultaneous application of several analysis methods (DEG analysis, KEGG analysis, K-mean cluster analysis, GO analysis, qPCR, network analysis, molecular docking involving dynamics simulation) allowed the expression patterns of genes and their functions to be revealed. The high transcription levels of genes encoding oxidoreductease and thiamine synthesis pathways can highlight and underline the knowledge on selection of promising candidate biocontrol agent. Additionally, the further studies on HOG1-thiamine interaction can extend our knowledge and encompass us to design synthetic compound mimicking thiamine, targeting on the specific side of the pathogen.

## 6 Author Contributions

RSS: Conceptualization, Data curation, Formal analysis, Funding acquisition, Investigation, Methodology, Project administration, Resources, Writing - original draft and Writing - review and editing. ÖB: Conceptualization, Data curation, Formal analysis, Investigation, Methodology, Resources, Software, Supervision, Validation, Visualization, Writing - original draft and Writing - review and editing. AC: Data curation, Formal analysis, Methodology. YK: Data curation, Formal analysis, Methodology. AK: Data curation, Formal analysis, Methodology. KKK: Data curation, Formal analysis, Methodology, Visualization. AÇ: Data curation, Formal analysis, Methodology. All authors contributed to the article and approved the submitted version.

## Acknowledgments

This study is dedicated to the 100th anniversary of the Republic of Turkiye. We are thankful to Assoc. Prof. Dr. Mustafa Küsek for Biolog Gen III analysis and Utku Berki Baysal (Junior computer engineer candidate) for Python coding support. The RNA-seq expression analysis of the whole raw data reads were also confirmed with Bacterial and Viral Bioinformatics Resource Center database (https://www.bv-brc.org) RNA-seq analysis pipeline.

## 9 Supplementary Material

The supplementary material for this article is accessible online in this preprint server.

## 12 Data Availability Statement

The datasets analyzed for this study can be found in the DNA Data Bank of Japan (http://trace.ddbj.nig.ac.jp/dra) under accession numbers SAMD00731600 (BioProject; PRJDB17356).

