## Supplementary material for "Exploring the genome-wide expression level of the bacterial strain belonging to *Bacillus safensis* (MM19) against *Phomopsis viticola*": Supplementary_Data_S1.docx

**Supplementary Data 1.** The sequences of 3 selected genes used for qPCR analysis.

**FUS3**

atgccgaacgcgcgcgtgtttaaaattctggcggcggcgaaactgaacaacattgcgctg

gaaattccggcgtatcagcatggcgtgaccaacaaaagcgcggaatttctgctgaaattt

ccggcgggcaaagtgccggcgtttgaaggcccggatggcttttgcctggtggaaagcgat

gcgattgcgcagtatgtggcgcagagcggcccgcaggcgagccagctgctgggccaggat

gcgatgagcagcgcgaaaattcgccagtggattagcttttttgcggaagaaatttatccg

accgtgctggatctggtgatgtggcgcgtgggcctgggcgcgtttgatgaaaccaccgaa

accaaagcgctgacccagctggtgtatggcctgagcgtgctggaaaaacatctgggcacc

ggcgcgctgctggtgggcgataaactgaccctggcggatctgaccggcgcgagcaccctg

ctgtgggcgtttatgcatattgtggatgaaccgatgcgccagcagtatccgaacgtggtg

gcgtggtatctgaaagtggtgcagaacgaagaagtggaagaagtgtttggcaaaccgaac

tttattgaaaaacgccgcctgggcgcgaaa

**qPCR primers**

**F-gaaaccaccgaaaccaaagc**

**R- gtgcccagatgtttttccag**

**HOG1**

atggcggaatttgtgcgcgcgcagatttttggcaccacctttgaaattaccagccgctat

agcgatctgcagccggtgggcatgggcgcgtttggcctggtgtgcagcgcgaaagataac

ctgaccggcagcaacgtggcggtgaaaaaaattatgaaaccgtttagcaccccggtgctg

agcaaacgcacctatcgcgaactgaaactgctgaaacatctgaaacatgaaaacgtgatt

agcctgagcgatatttttattagcccgctggaagatatttattttgtgaccgaactgctg

ggcaccgatctgcatcgcctgctgaccagccgcccgctggaaaaacagtttattcagtat

tttctgtatcagattctgcgcggcctgaaatatgtgcatagcgcgggcgtggtgcatcgc

gatctgaaaccgagcaacattctggtgaacgaaaactgcgatctgaaaatttgcgatttt

ggcctggcgcgcattcaggatccgcagatgaccggctatgtgagcacccgctattatcgc

gcgccggaaattatgctgacctggcagaaatatgatgtggaagtggatgtgtggagcgcg

ggctgcatttttgcggaaatgctggaaggcaaaccgctgtttccgggcaaagatcatgtg

aaccagtttagcattattaccgaactgctgggcaccccgccggatgatgtgattcatacc

attgcgagcgaaaacaccctgcgctttgtgcagagcctgccgaaacgcgaacgccagccg

ctggcgagcaaatttacccaggcggatccgctggcgattgatctgctggaaaaaatgctg

gtgtttgatccgcgcgcgcgcattaaagcggcggaaggcctggcgcatgaatatctgagc

ccgtatcatgatccgaccgatgaaccggcggcggaagaacgctttgattggagctttaac

gatgcggatctgccggtggatacctggaaaattatgatgtatagcgaaattctggattat

cataacgtgattaacgatgcgcagaacctgaccgaaagccag

**qPCR primers**

**F-attcaggatccgcagatgac**

**R-gccaggtcagcataatttcc**

**SGE1**

atggcgggcaccatgccgctgcgcccgacctatgtgggctttgtgcgcgataccaccgat

gcgctgctgatttttgaagcgtgcctgagcggcaccctgagccatgtgccgcgccgcccg

catgatcgcgaacgccaggatctgattaaaagcggcaacatttttgtgtatgaagaacat

gcgagcggcattaaacgctggaccgatagcattagctggagcccgagccgcattctgggc

aactatctgctgtatcgcgaactggaaaaaccgtttccgccgggcgaaaaaaaacgcgcg

cgcggccgcaacggcaaaagcaccacccagagcggcggcattagcaaagcgcgccagcgc

aacaccgtgccgtttccgcagggcctggaacatggcaacgaatatccgagcgtgccgagc

gatgatgaacgccatctggtgggcagcctggtggatagctatgattttaaagaacagggc

ctggtgaaaaaaaccattagcattacctatcagggcgtgccgcatcatctggtgagctat

tataacgtggaagatgtgaaagcgggcctgctgagcggcccgagcgatgatccgcgcctg

cgcggcgtggtgccgcgcaccgaactgatgaacggccagaactttcgcgcgccggtggaa

gaagcgatgggcggcagctatatgccgagcatggtggcgagcattggctatccgaccctg

cagcatcagagccagatgcatcagagccagatgcatcagccgcagatgcatcagccgcag

atgcatcagagccagatgcatcagagccagatgcatcagccgcagatgcatcagccgcag

gcgcatcagccgcaggtgcatcagccgcaggtgcatccgccgcaggtgcatcagccgcag

gcgcatcagccgcagtatcagagccagaccctgcatccgacccatggctatcagcagacc

tatgcgggccagccgaacgcgccgagcagcacctggtgg

**qPCR primers**

**F- ttaaacgctggaccgatagc**

**R- cagttcgcgatacagcagatag**

**Normalized genes**

| *AhActin-F* | CTGAAAGATTCCGATGCCCTGA |
| --- | --- |
| *AhActin-R* | AACCACCACTCAAGACAATGTTACCA |
