## Supplementary Figures for "Exploring the genome-wide expression level of the bacterial strain belonging to *Bacillus safensis* (MM19) against *Phomopsis viticola*": Supplementary_Figure_S1.docx

Supplementary Material

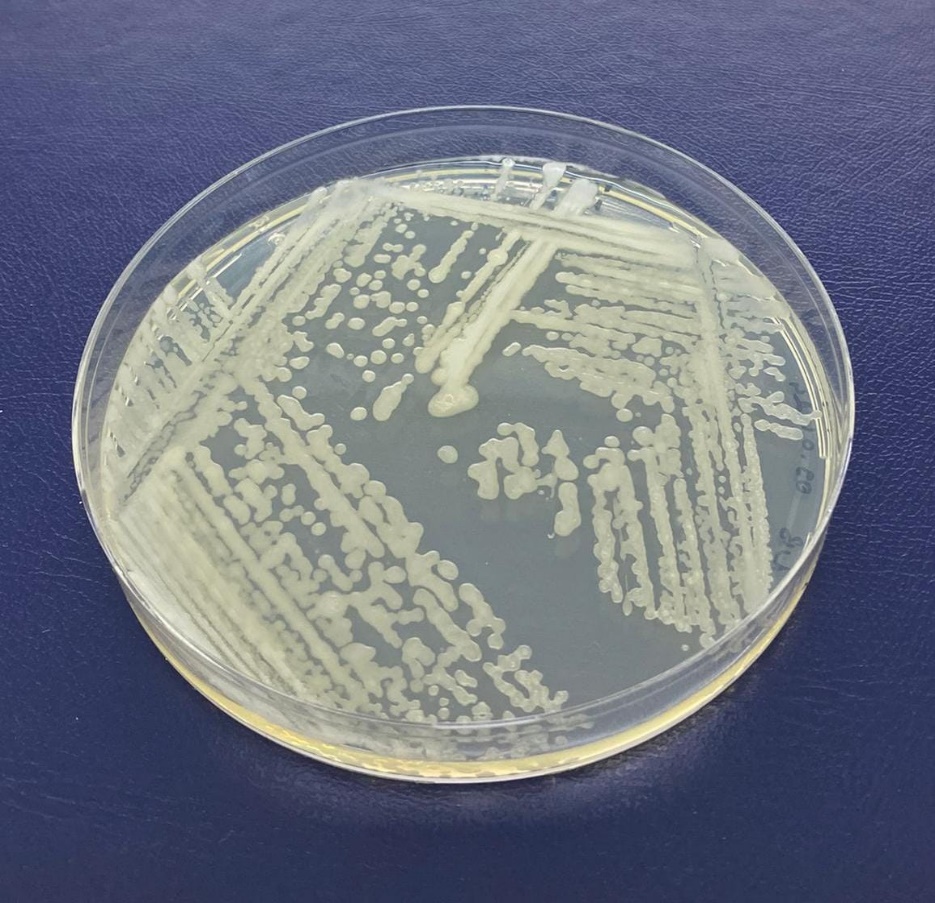

**Supplementary Figure S1.** Colony appearance of *Bacillus safensis* strain MM19 on Nutrient Agar.
