## Supplementary Figures for "Exploring the genome-wide expression level of the bacterial strain belonging to *Bacillus safensis* (MM19) against *Phomopsis viticola*": Supplementary_Figure_S2.docx

Supplementary Material

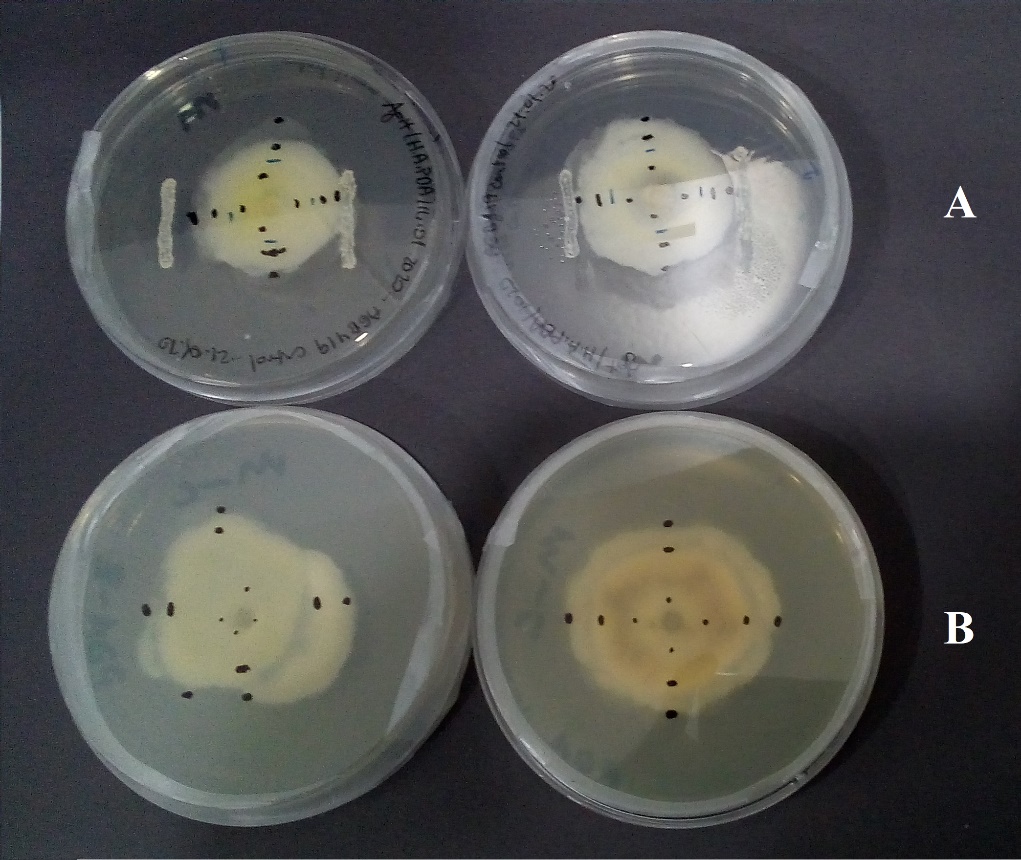

**Supplementary Figure S2.** *In vitro* dual culture assay. **(A)** bacteria-fungus interaction, **(B)** fungus control alone.
