## Supplementary Figures for "Exploring the genome-wide expression level of the bacterial strain belonging to *Bacillus safensis* (MM19) against *Phomopsis viticola*": Supplementary_Figure_S3.docx

Supplementary Material

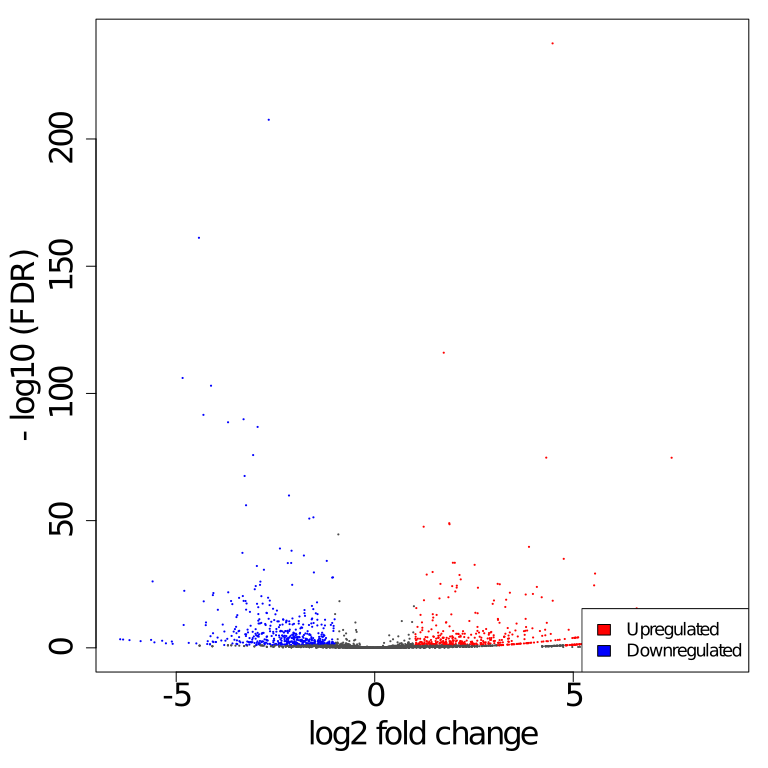

**Supplementary Figure S3.** Log2 fold change within up-regulated and down-regulated genes.
