## Supplementary Figures for "Exploring the genome-wide expression level of the bacterial strain belonging to *Bacillus safensis* (MM19) against *Phomopsis viticola*": Supplementary_Figure_S4.docx

Supplementary Material

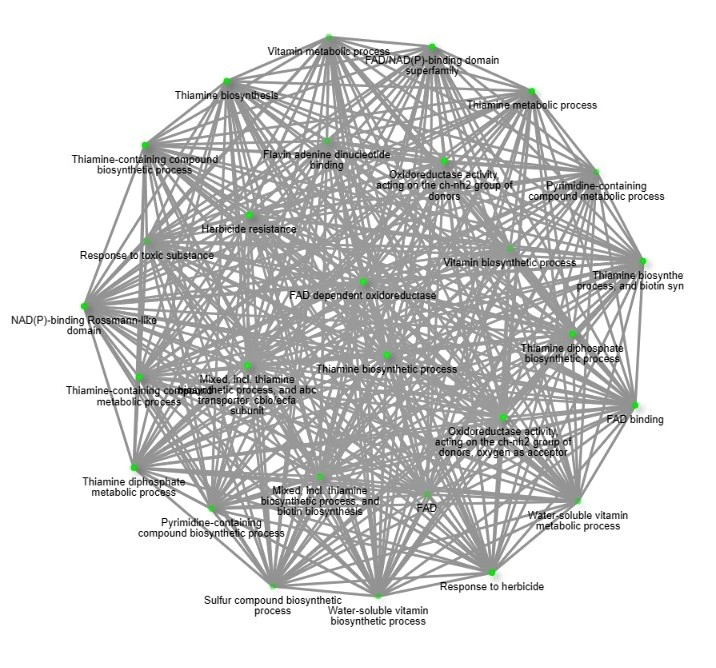

**Supplementary Figure S4.** Gene interaction analysis according to RNA-seq output.
