## Supplementary Figures for "Exploring the genome-wide expression level of the bacterial strain belonging to *Bacillus safensis* (MM19) against *Phomopsis viticola*": Supplementary_Figure_S5.docx

Supplementary Material

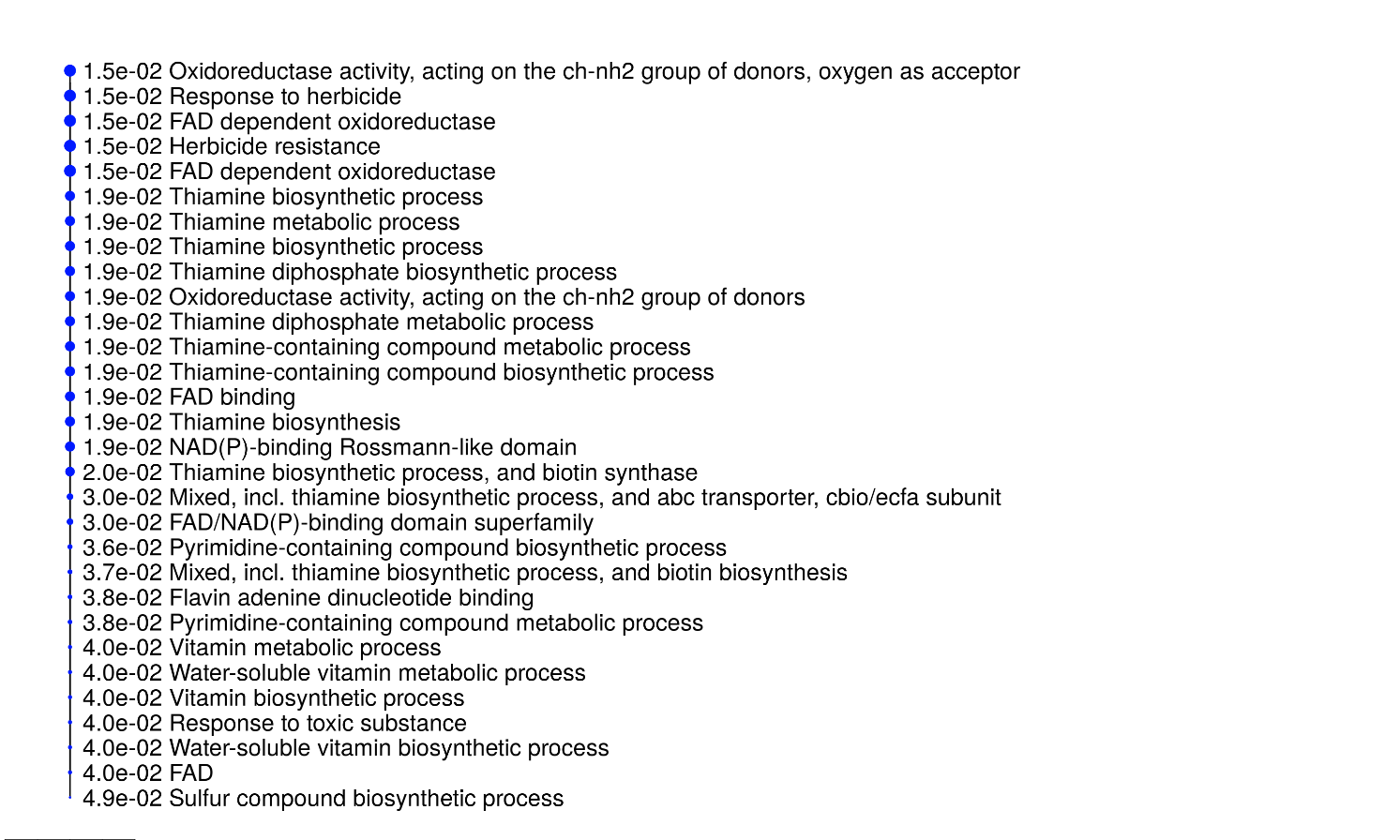

**Supplementary Figure S5.** Biological network analysis according to RNAseq output considering p values.
