## Supplementary Figures for "Exploring the genome-wide expression level of the bacterial strain belonging to *Bacillus safensis* (MM19) against *Phomopsis viticola*": Supplementary_Figure_S6.docx

Supplementary Material

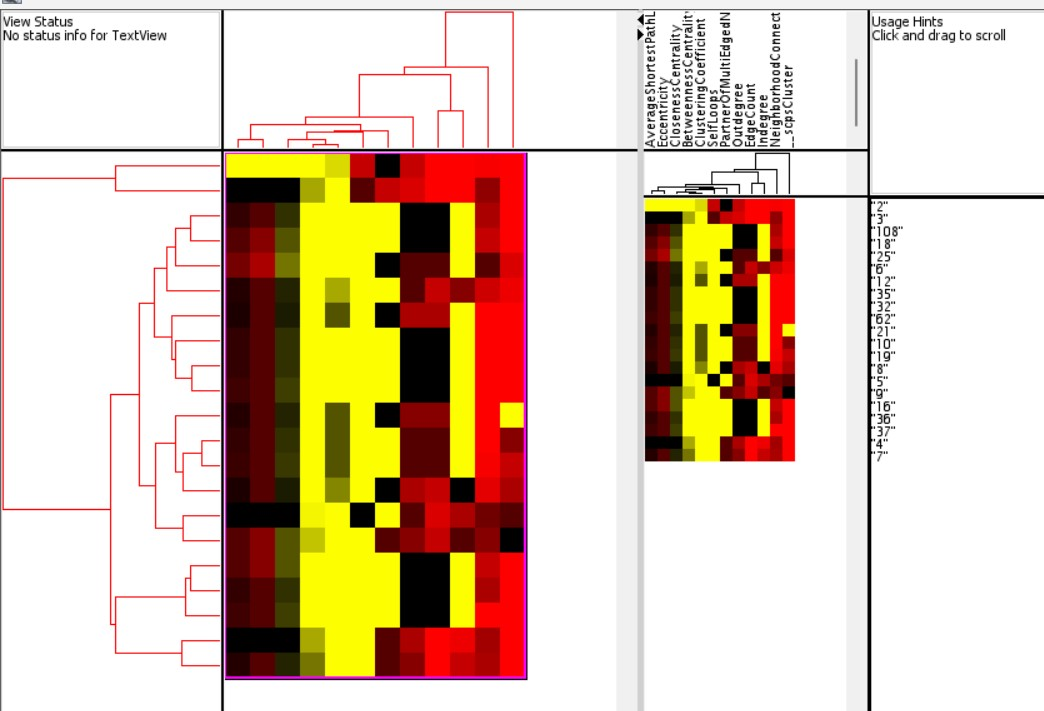

**Supplementary Figure S6.** Cytoscape gene network tree analysis according to RNA-seq output considering most variable top 20 genes in bacteria during the fungi-bacteria interaction.
