## Supplementary Tables for "Exploring the genome-wide expression level of the bacterial strain belonging to *Bacillus safensis* (MM19) against *Phomopsis viticola*": Supplementary_Table_S1.docx

Supplementary Material

| **Utilisation of carbohydrates** | **Results** | **Utilisation of amino acids and derivatives** | **Results** | **Utilisation of carboxylic acids and derivatives** | **Results** | **Resistance to chemicals** | **Results** |
| --- | --- | --- | --- | --- | --- | --- | --- |
| Dextrin | + | L-alanine | + | Citric acid | + | pH 7 | + |
| D-trehalose | + | L-glutamic acid | + | L-malic acid | + | pH 6 | + |
| β-methyl-D-glucoside | + | L-aspartic acid | + | Acetoacetic acid | + | pH 5 | + |
| α-D-glucose | + | L-histidine | - | L-lactic acid | + | 1% NaCl | + |
| D-cellobiose | + | L-serine | - | Acetic acid | - | 4% NaC | + |
| Gentiobiose | + | Glycyl-L-proline | - | Formic acid | + | 8% NaCl | + |
| Sucrose | + | Gelatin | - | Methyl pyruvate | - | 1% sodium  lactate | + |
| D-fructose | + | D-serine | - | Bromo-succinic-acid | - | Lithium chloride | + |
| D-maltose | + | D-aspartic acid | - | Tween 40 | - | Potassium tellurite | + |
| D-mannose | + | L-arginine | - | β-hydroxy-D,L-butyric acid | - | Guanidine HCl | - |
| N-acetyl-D-glucosamine | - | L-pyroglutamic  acid | - | γ-aminobutryric acid | - | Aztreonam | + |
| D-salicin | + |  |  | Propionic acid | - | Sodium butyrate | + |
| D-raffinose | + |  |  | ρ-hydroxy phenylacetic acid | - | Rifamycin SV | + |
| D-galactose | - |  |  | α-keto-glutaric-acid | - | Sodium bromate | - |
| Inosine | - |  |  | D-malic acid | - | Tetrazolium  violet | - |
| D-turanose | + |  |  | α-hydroxy-butyric acid | - | Tetrazolium-blue | - |
| Stachyose | - |  |  | D-lactic acid-methyl ester |  | Troleandomycin | - |
| D-melibiose | - |  |  | α-keto-butyric-acid | - | Lincomycin | - |
| N-acetyl-β-D-mannosamine | - |  |  |  |  | Niaproof 4 | - |
| α-D-lactose | - |  |  |  |  | Vancomycin | - |
| L-rhamnose | - |  |  |  |  | Fusidic acid | - |
| N-acetyl-D-galactosamine | - |  |  |  |  | Minocycline | - |
| α-D-lactose | - |  |  |  |  | Nalidixic acid | - |
| L-rhamnose | - |  |  |  |  |  |  |
| N-acetyl-D-galactosamine | - |  |  |  |  |  |  |
| N-acetyl neuraminic acid | - |  |  |  |  |  |  |
| D-fucose | - |  |  |  |  |  |  |
| 3-methyl glucose | - |  |  |  |  |  |  |
| L-fucose | - |  |  |  |  |  |  |
| Glycerol | + |  |  |  |  |  |  |
| D-mannitol | + |  |  |  |  |  |  |
| D-sorbitol | + |  |  |  |  |  |  |
| Myo-inositol | - |  |  |  |  |  |  |
| D-arabitol | - |  |  |  |  |  |  |
| D-fructose-6-phosphate | + |  |  |  |  |  |  |
| D-glucose-6-phosphate | - |  |  |  |  |  |  |
| Pectin | + |  |  |  |  |  |  |
| Glucuronamide | + |  |  |  |  |  |  |
| D-gluconic acid | + |  |  |  |  |  |  |
| D-galacturonic acid | + |  |  |  |  |  |  |
| L-galactonic-acid lactone | + |  |  |  |  |  |  |
| D-glucuronic-acid | + |  |  |  |  |  |  |
| D-saccharic acid | + |  |  |  |  |  |  |
| Mucic acid | + |  |  |  |  |  |  |
| Quinic acid | - |  |  |  |  |  |  |

**Supplementary Table S1.** The phenotypic profile of *Bacillus safensis* strain MM19 as determined by Biolog GENIII MicroPlate assay. +, Positive utilization and −, negative utilization.
