## Supplementary Tables for "Exploring the genome-wide expression level of the bacterial strain belonging to *Bacillus safensis* (MM19) against *Phomopsis viticola*": Supplementary_Table_S2.docx

***Supplementary Material***

**Supplementary Table S2.** T-test analysis on inhibitory effect of *Bacillus safensis* strain *MM19* against *Phomopsis viticola* *in vitro*.

**T-Test**

| **Notes** | | |
| --- | --- | --- |
| Output Created | | 18-OCT-2023 14:55:44 |
| Comments | |  |
| Input | Active Dataset | DataSet0 |
|  | Filter | <none> |
|  | Weight | <none> |
|  | Split File | <none> |
|  | N of Rows in Working Data File | 7780 |
| Missing Value Handling | Definition of Missing | User defined missing values are treated as missing. |
|  | Cases Used | Statistics for each analysis are based on the cases with no missing or out-of-range data for any variable in the analysis. |
| Syntax | | T-TEST GROUPS=Treatment(1 2)  /MISSING=ANALYSIS  /VARIABLES=D7 D10 D12  /CRITERIA=CI(.95). |
| Resources | Processor Time | 00:00:00.36 |
|  | Elapsed Time | 00:00:03.14 |

| **Group Statistics** | | | | | | | |
| --- | --- | --- | --- | --- | --- | --- | --- |
|  | **Treatment** | | **Statistic** | **Bootstrap^a^** | | | |
|  |  |  |  | **Bias** | **Std. Error** | **95% Confidence Interval** | |
|  |  |  |  |  |  | **Lower** | **Upper** |
| D7 | Control | N | 6 |  |  |  |  |
|  |  | Mean | 1.9333 | .0014 | .0529 | 1.8400 | 2.0500 |
|  |  | Std. Deviation | .12910 | **-.01657^b^** | .03814^b^ | **.02672^b^** | **.17536^b^** |
|  |  | Std. Error Mean | .05270 |  |  |  |  |
|  | Treated | N | 6 |  |  |  |  |
|  |  | Mean | 1.5333 | -.0007 | .0493 | 1.4376 | 1.6300 |
|  |  | Std. Deviation | .12910 | **-.02105^c^** | .04362^c^ | **.02440^c^** | **.19149^c^** |
|  |  | Std. Error Mean | .05270 |  |  |  |  |
| D10 | Control | N | 6 |  |  |  |  |
|  |  | Mean | 3.4417 | .0006 | .0253 | 3.3929 | 3.5000 |
|  |  | Std. Deviation | .06646 | **-.00897^b^** | .02122^b^ | **.00000^b^** | **.09574^b^** |
|  |  | Std. Error Mean | .02713 |  |  |  |  |
|  | Treated | N | 6 |  |  |  |  |
|  |  | Mean | 2.7000 | .0020 | .1599 | 2.3667 | 3.0061 |
|  |  | Std. Deviation | .41231 | **-.05184^c^** | .10799^c^ | **.11180^c^** | **.53947^c^** |
|  |  | Std. Error Mean | .16833 |  |  |  |  |
| D12 | Control | N | 6 |  |  |  |  |
|  |  | Mean | 4.8417 | .0001 | .0649 | 4.7083 | 4.9700 |
|  |  | Std. Deviation | .16857 | **-.02045^b^** | .04109^b^ | **.05000^b^** | **.21361^b^** |
|  |  | Std. Error Mean | .06882 |  |  |  |  |
|  | Treated | N | 6 |  |  |  |  |
|  |  | Mean | 3.9417 | .0072 | .2300 | 3.4750 | 4.3749 |
|  |  | Std. Deviation | .58174 | **-.08007^c^** | .17591^c^ | **.15275^c^** | **.82702^c^** |
|  |  | Std. Error Mean | .23749 |  |  |  |  |
| a. Unless otherwise noted, bootstrap results are based on 1000 bootstrap samples | | | | | | | |
| b. Based on 998 samples | | | | | | | |
| c. Based on 999 samples | | | | | | | |

| **Independent Samples Test** | | | | | | | | | | |
| --- | --- | --- | --- | --- | --- | --- | --- | --- | --- | --- |
|  | | **Levene's Test for Equality of Variances** | | **t-test for Equality of Means** | | | | | | |
|  |  | **F** | **Sig.** | **t** | **df** | **Sig. (2-tailed)** | **Mean Difference** | **Std. Error Difference** | **95% Confidence Interval of the Difference** | |
|  |  |  |  |  |  |  |  |  | **Lower** | **Upper** |
| D7 | Equal variances assumed | .128 | .728 | 5.367 | 10 | .000 | .40000 | .07454 | .23392 | .56608 |
|  | Equal variances not assumed |  |  | 5.367 | 10.000 | .000 | .40000 | .07454 | .23392 | .56608 |
| D10 | Equal variances assumed | 12.935 | .005 | 4.350 | 10 | .001 | .74167 | .17050 | .36177 | 1.12156 |
|  | Equal variances not assumed |  |  | 4.350 | 5.260 | .007 | .74167 | .17050 | .30982 | 1.17352 |
| D12 | Equal variances assumed | 2.991 | .114 | 3.640 | 10 | .005 | .90000 | .24726 | .34906 | 1.45094 |
|  | Equal variances not assumed |  |  | 3.640 | 5.834 | .011 | .90000 | .24726 | .29077 | 1.50923 |

| **Bootstrap for Independent Samples Test** | | | | | | | |
| --- | --- | --- | --- | --- | --- | --- | --- |
|  | | **Mean Difference** | **Bootstrap^a^** | | | | |
|  |  |  | **Bias** | **Std. Error** | **Sig. (2-tailed)** | **95% Confidence Interval** | |
|  |  |  |  |  |  | **Lower** | **Upper** |
| D7 | Equal variances assumed | .40000 | .00205 | .07134 | .001 | .26671 | .55000 |
|  | Equal variances not assumed | .40000 | .00205 | .07134 |  | .26671 | .55000 |
| D10 | Equal variances assumed | .74167 | -.00134 | .16168 | .042 | .44010 | 1.07493 |
|  | Equal variances not assumed | .74167 | -.00134 | .16168 |  | .44010 | 1.07493 |
| D12 | Equal variances assumed | .90000 | -.00707 | .24190 | .011 | .41951 | 1.36875 |
|  | Equal variances not assumed | .90000 | -.00707 | .24190 |  | .41951 | 1.36875 |
| a. Unless otherwise noted, bootstrap results are based on 1000 bootstrap samples | | | | | | | |
